# Evolution of genotypic and phenotypic diversity in multispecies biofilms

**DOI:** 10.1101/2023.10.08.561388

**Authors:** Cristina I. Amador, Sofia Zoe Moscovitz, Lorrie Maccario, Jakob Herschend, Isabel-Sophie Kramer, Hannah Jeckel, Vaughn S. Cooper, Knut Drescher, Thomas R. Neu, Mette Burmølle, Henriette L. Røder

## Abstract

Bacterial fitness and adaptability in microbial communities are influenced by interspecies interactions and spatial organization. This study investigated how these factors shape the evolutionary dynamics of *Bacillus thuringiensis*. A distinct phenotypic variant of *B. thuringiensis* emerged consistently under both planktonic and biofilm conditions, as well as in monospecies and mixed-species settings, but was strongly selected in biofilms and during coexistence with *Pseudomonas defluvii* and/ or *Pseudomonas brenneri*. Compared to its ancestor, the variant exhibited shorter generation times, reduced sporulation, auto-aggregation, and lower biomass in mixed-species biofilms. Mutations in the *spo0A* regulator, which controls sporulation and biofilm matrix production, were identified in all variants. Proteomics revealed a reduction in TasA, a key matrix protein, in the variant but increased levels in co-culture with *P. brenneri*. These findings highlight how interspecies interactions drive *B. thuringiensis* diversification, promoting traits like reduced matrix production and species coexistence, with implications for microbial consortia applications in agriculture and biopesticides.

## Introduction

When encountering new environmental challenges, bacterial populations can adapt their physiological state (phenotype) through different mechanisms: expressing specific traits, acquiring new advantageous mutations (*de novo* genetic variation), or utilizing the pre-existing genetic diversity within the population (standing genetic variation) [1–3]. Besides intraspecific variation, natural communities contain many species that might either constrain adaptation by competing for resources, e.g., via niche exclusion [4], or facilitate adaptation by making new niches available [5, 6]. In biofilms, microbial cells are in close proximity and can influence each otheŕs phenotypes and evolutionary trajectories through social interactions [7]. Spatial segregation in biofilms increases the frequency of interactions between cells of the same genotype and favors cooperative behaviors but can also support variants that are otherwise outcompeted in unsegregated environments [8–10] and facilitate coexisting populations [11, 12]. Several studies have reported that competitive interactions within biofilms are dominant, including competition for limited space and resources [10, 13–16]. However, evidence shows that competition can subside locally over time, often leading to stable coexistence [17]. We have previously described that interspecies interactions can lead to evolved variants that display enhanced mutualism in biofilms [18–20]. Thus, to better understand bacterial adaptation mechanisms in natural environments, investigations of adaptation mechanisms in the presence of spatial structure and multiple species that co-occur in natural communities are crucial.

The evolution of social interactions within biofilms has been extensively studied in various Gram-negative bacteria, including *Pseudomonas spp*. and *Burkholderia cenocepacia* [21–23]. However, comparatively less attention has been devoted to investigating evolutionary processes of Gram-positive bacteria, especially in the presence of multiple species. While both bacterial groups may be subjected to similar evolutionary forces, they possess distinct biological and ecological features, which may lead to divergent adaptive dynamics. Although some studies have explored the adaptive evolution of Gram-positive bacteria in diverse biofilm systems, their focus has primarily been on adaptive genotypes [24–26] or co-culture [20, 27]. Such limited focus leaves a gap in our understanding of how Gram-positive bacteria evolve within complex, multispecies biofilms.

In this study, we aim to investigate how cultivation conditions—specifically spatial structure (biofilm *vs.* planktonic settings) and interspecies interactions (mono-, dual- or multispecies)— influence the diversification of *Bacillus thuringiensis* (BT), a Gram-positive, spore-forming bacterium renowned for its insecticidal capabilities [28]. BT was cultured in isolation and in combination with *Pseudomonas defluvii* (PD) and *Pseudomonas brenneri* (PB), both of which were co-isolated with BT from a wastewater treatment facility in Kalundborg, Denmark [29]. *Bacillus* spp. and *Pseudomonas* spp. are known for their utility as plant growth enhancers and biopesticides [30]. Importantly, research has demonstrated that such bacterial consortia manifest unique, emergent properties not attainable by individual species alone [31]. Additionally, the rationale behind selecting these species for our investigation stemmed from their potential synergistic contributions to biofilm formation, a premise supported by preliminary laboratory findings. Other studies on laboratory adaptive evolution employed biofilm settings focusing on biofilm dispersal and recolonization over short evolutionary spans [20, 32]. However, our work aimed at investigating the dynamics of cells that remained in the biofilm, attached to plastic slides. This led to the identification of a phenotypic variant of BT, termed “light variant,” from its reduced Congo red dye binding in colonies on agar, compared to the ancestral strain. Morphotypic colony diversification often reflects underlying genotypic changes, which play a crucial role in shaping biofilm formation and adaptation [18, 33, 34]. Therefore, we isolated and characterized such light variants, which arose frequently. We found that the light variant colony morphotype was linked to mutations in sporulation genes, and that these variants demonstrated a remarkable ability to dominate and replace the wild-type phenotype in most conditions examined. Crucially, the fitness of this evolved phenotype was enhanced in mixed-species cultures and within biofilm contexts, underlining the effects of both biotic and abiotic complexity on evolutionary dynamics. This finding sheds light on the adaptive strategies of BT within microbial consortia.

## Results

### Emergence of a BT phenotypic variant after short-term evolution assay

We investigated how the presence of other species influenced the selection of new phenotypic variants of BT under biofilm or planktonic conditions. For the biofilm condition, cells were incubated in TSB with a submerged polycarbonate slide. Prior to transferring the slides, we removed and discarded the floating pellicle, when present, since our evolution experiment targeted surface-adhered cells. The polycarbonate slides were then removed from the well and washed with PBS to remove unattached cells, and transferred to new wells containing fresh media, followed by incubation for another 24 h. This process was repeated for eight consecutive cycles. On day 8, we plated and enumerated the cultures based on distinct colony morphology (Figure 1a). Although not all genetic changes manifest in noticeable physical properties, genetic variation can drive morphotypic colony variation, especially in biofilm settings [18, 33, 34]. In this study, we focused on selecting variants of BT forming colonies of varying morphotypes, by employing TSA Congo Red. Congo red binds different biofilm matrix components, including amyloids and polysaccharides [35, 36]. Thus, altered colony morphology on Congo red agar could provide a visual marker of the underlying genetic changes.

**Figure 1.**
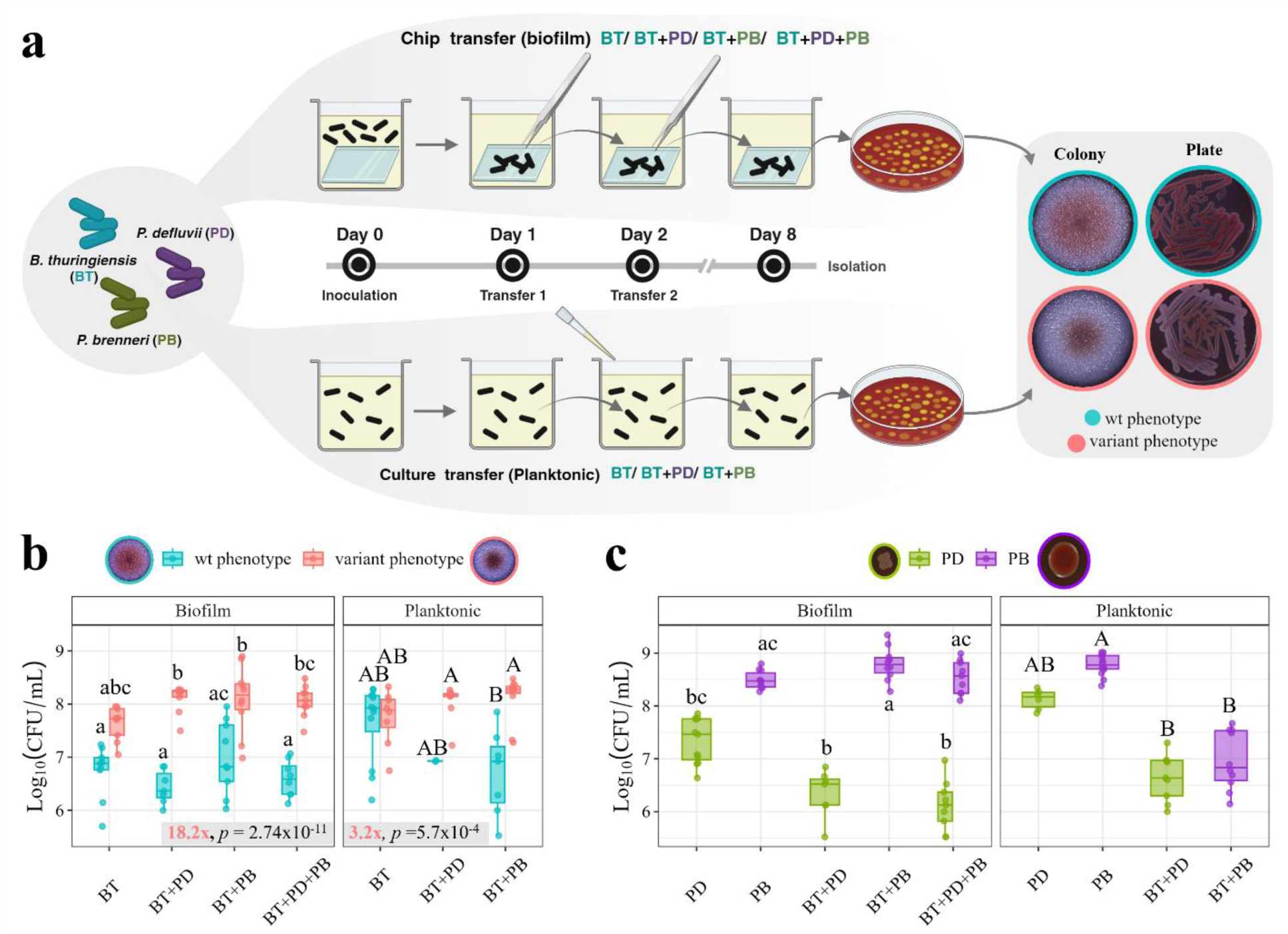
Short-term evolution assay of *Bacillus thuringiensis* (BT) in the presence/absence of *Pseudomonas* species. **(a)** Experimental setup: BT was cultured alone or with *Pseudomonas brenneri* (PB) and/or *Pseudomonas defluvii* (PD) in Tryptic Soy Broth (TSB) medium, either as planktonic or biofilm cultures, for 24 hours. Biofilm samples were transferred daily onto fresh medium for eight consecutive days, followed by plating on day 8 and CFU counting on TSB agar supplemented with 40 µg/mL Congo red and 20 µg/mL Coomassie brilliant blue (G). **(b)** BT cell counts (Log_10_(CFU/mL)) of variant (red) and wild-type (blue) phenotypes in different strain combinations after the evolution experiment, presented for biofilm (left) and planktonic (right) cultures. Box plots represent ≥ 3 biological replicates, represented as individual points, each averaged from 3 technical replicates. Different letters denote significance per facet (Dunńs post-hoc test, *p* < 0.05, Bonferroni correction). Gray boxes represent CFU/mL ratios of variant *vs*. wild-type phenotypes within each setting (biofilm or planktonic) and statistical significance of such ratios (Wilcoxon’s test comparing variant and wild-type phenotypes per setting, *p* < 0.05). **(c)** Cell counts (Log_10_(CFU/mL)) of PD (green) and PB (purple) in various strain combinations and culture conditions after the evolution experiment. Box plots represent ≥ 7 biological replicates, represented as in (b). Different letters denote significance per facet (Dunńs post-hoc test, *p* < 0.05, Bonferroni correction).

A particular BT phenotypic variant was detected in all replicate evolution experiments, which we termed “light variant” as it bound less Congo red compared to the ancestor (Figure 1a). Colonies exhibiting phenotypic traits indistinguishable from those of the ancestral BT strain— derived from the evolution experiment—are referred to as wild-type phenotype throughout this study. When comparing within colony phenotype depending on the culture condition (wild-type mono-culture *vs.* co-culture; or variant mono-culture *vs.* co-culture), no significant differences in population size were detected (*p* > 0.05, Dunn’s post-hoc test; Figure 1b and Supplementary Table 1). However, when comparing BT wild-type and light variant colony phenotypes per culture condition, the latter emerged independently in each evolved biofilm population where it consistently grew to significantly higher CFU counts than the wild-type phenotype (*p* < 0.05, Dunn’s post-hoc test; Figure 1b and Supplementary Table 1).

On the contrary, analogous evolution experiments in planktonic culture only showed significant difference in CFU counts between the wild-type and light variant phenotypes in co-culture with PB (*p* = 5.07×10^-04,^ Dunn’s post-hoc test; Figure 1b and Supplementary Table 1). Still, the variant phenotype was found to be significantly more prevalent than the wild-type phenotype regardless of the cultivation settings— biofilm or planktonic— when not considering the species combination (Wilcoxon’s test, *p*_Biofilm_ = 2.74×10^-11^; *p*_Plank_ = 5.7×10^-05^, Figure 1b). However, the CFU/ mL ratio of variant to wild-type phenotype was substantially higher in biofilm compared to planktonic settings (Ratio variant-wt: biofilm = 18.2-fold; planktonic = 3.2-fold, Figure 1b). To ensure statistical robustness, we re-analyzed the data excluding the BT + PD + PB treatment from the biofilm condition. Nonetheless, the variant still emerged at significantly higher frequency than the wild-type phenotype under biofilm conditions (Ratio variant-wt = 12.2-fold, *p*_Biofilm_ = 2.32×10^-08^, Wilcoxon’s test), supporting the conclusion that biofilm evolution enhances variant emergence. Hence, our results from short-term evolution experiments suggest that the BT light variant exhibited higher fitness under biofilm conditions and in the presence of PB and/or PD when compared to the wild-type colony phenotype.

During the evolution experiment, all species were maintained. However, the trajectories of the *Pseudomonas* species varied depending on the culture condition and design, as shown in Figure 1c. BT showed a competitive interaction with both *Pseudomonas* species in planktonic settings (Figure 1c, right panel). Specifically, the presence of BT reduced the number of PB cells by 1.92 Log_10_ (Dunn’s post-hoc test; *p* = 2.7×10^-05^, Log_10_ CFU/mL: BT+PB = 6.88; PB = 8.8). These findings indicate that the lack of spatial structure and resulting intermixing may limit the co-existence of different species. However, this was not observed under biofilm conditions, where no significant differences were observed in *Pseudomonas* populations, despite the presence/ absence of BT (*p* = 1.0, Dunn’s post-hoc test; Figure 1c). This suggests that the adapted BT phenotype may facilitate the co-existence of BT and PB in biofilm settings.

### The BT “light variant” exhibited distinct phenotypic traits compared to its ancestor

We isolated 150 BT colonies of both light and wild-type phenotype (50% of each) from independent evolution experiments and conditions. All colonies within each phenotype group (light or wild type) displayed similar phenotypic traits. The BT light variant displayed different colony morphology compared to the ancestral BT strain, because of reduced Congo red binding (Figure 1a). Congo red binds to different biofilm matrix components, including amyloids and polysaccharides [35, 36]. Furthermore, the wild-type and light variant phenotypes also had visibly different phenotypes during growth in planktonic cultures: under non-shaking conditions, the ancestral BT cells sank to the bottom of the culture, while cells of the light variant remained in suspension (Figure 2a). Auto-aggregation, where bacteria attach to each other rather than to a surface, is observed in many bacterial species. This behavior involves the formation of macroscopic cell aggregates that settle at the bottom of culture tubes. [37, 38]. Moreover, some environmental *Bacillus licheniformis* have been documented to undergo aggregation during suspension culture under carbon limitation or glucose-free conditions [39, 40].

**Figure 2.**
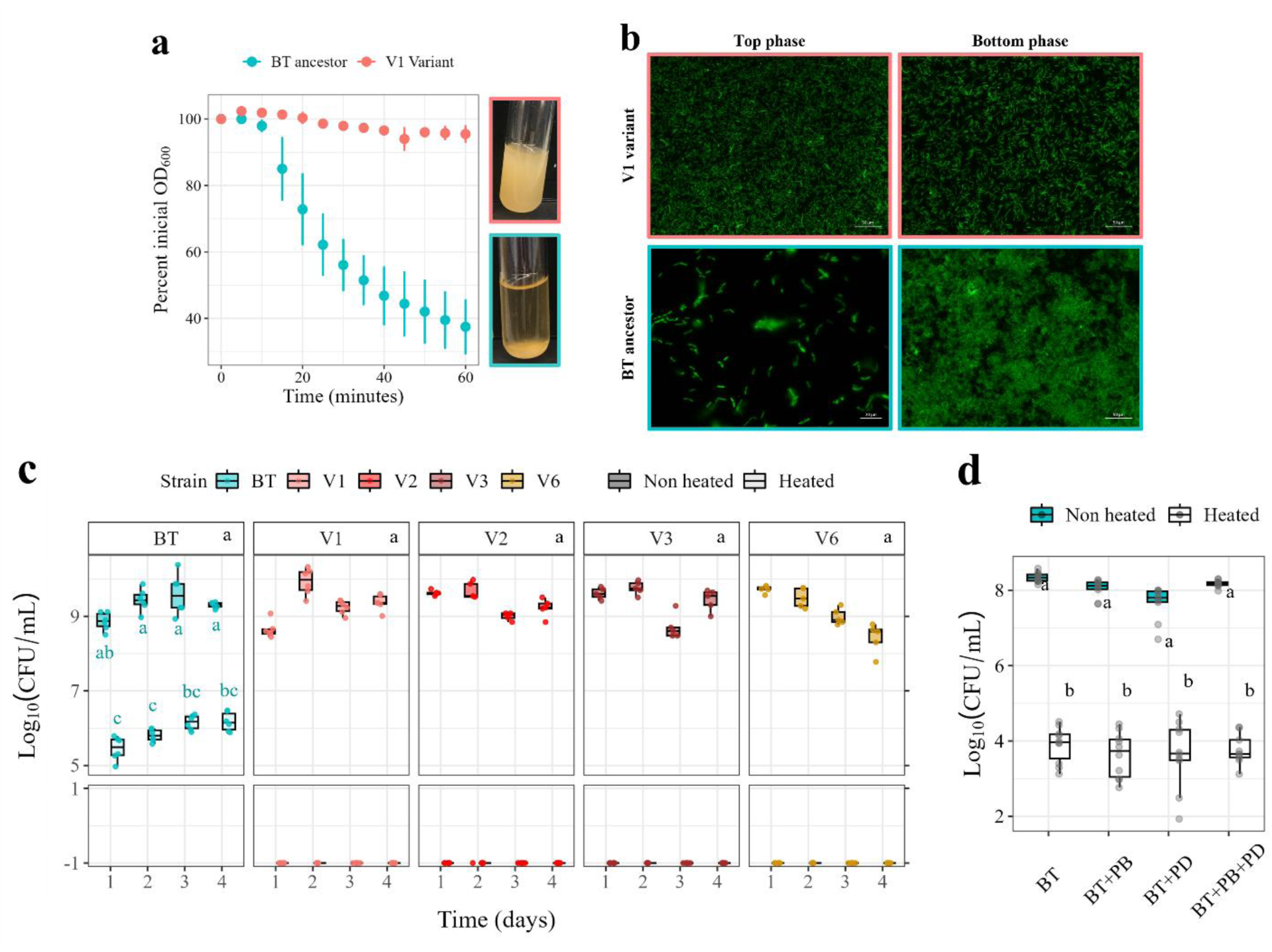
Phenotypic features of *B. thuringiensis* ancestor and adapted variant strains. **(a**) Sedimentation pattern of BT ancestor and V1 variant strains in TSB planktonic cultures. Sedimentation dynamics was measured as the decay of initial OD_600_ over time, represented as the mean OD_600_ percentage relative to initial OD_600_ (t_0_). Data based on 3 biological replicates. In the absence of shaking, sedimentation can be observed only for the ancestor strain. **(b**) Auto-aggregative phenotype of BT ancestral strain compared to the adapted variant. Images show top and bottom phases of planktonic cultures in TSB stained with 10 µM SYTO 16 nucleic acid cell stain. Images were acquired with a Zeiss Axio Observer Z1 epifluorescence microscope with a LD-NEOFLUAR 20x objective. Scale bars represent 50 µm. **(c)** Spore formation ability of *B. thuringiensis* ancestral (BT) and variant strains under control (vegetative cells, non-heated) and boiling (germinating spores, heated) conditions in planktonic monocultures over 1-4 days incubation. Log_10_(CFU/mL) values were adjusted by adding 0.1 to accommodate zero values (represented as −1). n ≥ 5 biological replicates. Different letters denote significant differences (*p* < 0.05, Dunńs post-hoc test, Bonferroni correction). Black letters denote significant differences between strains only considering non-heated conditions, while turquoise letters indicate comparisons within BT (heated *vs.* non-heated). **(d)** Spore formation ability of BT ancestral strain in biofilm settings in different strain combinations after one-day incubation; n ≥10 biological replicates.

We quantified aggregation by measuring the percentage of initial OD_600_ remaining in suspension over time, using the variant and ancestral BT strains. The variant cells remained mainly in suspension (95%), whereas the ancestral BT strain showed higher cell sedimentation, retaining only 37% of its initial OD_600_ (Figure 2a). Additionally, we also investigated aggregation in both strains by microscopic analysis. For this, we stained top and bottom phases of planktonic cultures for both strains with the nucleic acid stain SYTO 16. We found that the ancestral BT strain predominantly formed cell clusters in the bottom phase, whereas the variant consisted mainly of single cells regardless of the visualized phase (Figure 2b). Thus, we confirm that the observed sedimentation by the BT ancestor is linked to auto-aggregation, whereas cells of the variant phenotype do not aggregate to the same extent.

In addition to these phenotypic traits, the BT light variant was unable to produce spores unlike the ancestral BT strain, as evidenced by optical microscopy (not shown). Given that all variant phenotype colonies displayed similar phenotypic traits, we focused on four light variants, namely V1, V2, V3, and V6, and further investigated their characteristics. These light variants were selected from different culture conditions and evolutionary lineages (independent evolutionary experiments): V1 and V2 were isolated from biofilm co-cultures with PB in different lineages (B and C, respectively), whereas V3 and V6 were isolated from BT biofilm mono-cultures (lineages B and C, respectively) (Supplementary Table 2).

We measured the capacity for spore formation of both ancestral and variant strains in planktonic settings over four days. These assays demonstrated that the variants did not produce spores, whereas the ancestral BT strain consistently produced spores regardless of the incubation time (Figure 2c). Next, we investigated whether other species may influence spore formation in the ancestor. The ability to form spores was assessed in mono-, two-species-, or multispecies biofilm conditions for 24-hour biofilms; however, no significant difference was observed between the tested combinations (Dunńs post-hoc test, *p* > 0.05; Figure 2d).

We hypothesized that selection for faster growth may have favored the loss of spore production in BT light variants, as sporulation is an energy-intensive process. Growth curves were performed for the BT ancestral and light variant strain in TSB. We found that the four BT light variants consistently showed a significantly faster doubling time (t_gen_) and growth rate (r) compared to their ancestor (Supplementary Table 3). Consequently, the time required to reach half of the maximum culture yield was significantly lower (t_mid_ or ½ k). This data suggests that the BT light variants have a growth advantage compared to the ancestral BT when competing for resources with the *Pseudomonas* species. This may also have led to the higher CFU counts observed in mixed-species cultures during our short-term evolution experiments in biofilms and planktonic cultures.

### BT light variants displayed mutations in sporulation-related genes

Whole genome sequencing of BT light variants and the BT ancestor revealed that the light variants displayed mutations in sporulation-related genes. This correlated with the observed lack of sporulation in all variants. Variants V1, V3, and V6 contained distinct mutations resulting in loss of function of the *spo0A* gene, as shown in Supplementary Figure 1a). The *spo0A* gene encodes a response regulator that, in its phosphorylated active form, activates the transcription of early sporulation genes and, consequently, the initiation of sporulation (Supplementary Figure 1b) [41, 42]. For variants V1 and V3, large deletions were observed in this region. Specifically, V1 lacked a 1.5 kb fragment containing most of *spo0A* and CDS0080 genes, whereas V3 had a 4.5 kb deletion that also included the *spoIVB* and *recN* genes (Supplementary Figure 1a). V6 only contained a non-synonymous mutation in the *spo0A* gene (C151>T), resulting in a change in the amino acid sequence Q51>stop, leading to early termination and, thus, loss of function of Spo0A. These mutations, confirmed by Sanger sequencing using specific primers (data not shown), undoubtedly explain the non-sporulating phenotype for variants V1, V3, and V6, given the central role of *spo0A* and *spoIVB* in the sporulation process. In regards to variant V2, we didńt find mutations in *spo0A* but structural variation in the *soj* gene, that is, large-scale changes in the structure of DNA [43]. Soj exerts a dual role in chromosome partitioning and sporulation, particularly in regulating chromosome dynamics as the cell undergoes significant structural changes [44]. All the mutations found in the variants are listed in Supplementary Table 4.

Additionally, variant V1 lacked 83 Kb region, including 40 hypothetical proteins and 6 IS family transposases, among other genes. However, none of the genes comprised in such region seemed to be related to sporulation (Supplementary Table 4). Furthermore, this variant had identical phenotypic traits to all other selected variants.

### Reduced biofilm matrix production in mixed biofilms with the adapted variant

The BT light variant phenotype showed increased CFU counts compared to the wild-type phenotype after the eight-day evolution experiment, especially in biofilm settings (adhesion to slides). However, the ancestral strain showed auto-aggregative phenotype in liquid cultures, unlike the variants, suggesting a different pattern for the formation of cell aggregates (non-associated to a surface). We thus wondered whether population size in biofilm settings may correlate with biofilm formation of the evolved phenotype. We measured the total amount of biofilm formed (including cells and matrix) on the slides after 24 hours by quantification of crystal violet (CV) retention as an overall estimate of biofilm biomass. We also quantified the specific prevalence of each strain by CFU counts of both the BT ancestral and V1 variant strains (as shown in Figure 3). Biofilm biomass was similar for the BT ancestor and variant in mono-culture while co-cultivation of ancestor and variant strains with PB, or in multispecies conditions, induced biofilm biomass in both. However, when comparing mono-culture *vs.* mixed-culture conditions, such biomass induction was only significant for the ancestral strain (Dunńs post-hoc test, BT *vs.* BT+PB, *p* = 0.0021; BT *vs.* BT+PD+PB, *p* = 0.0106; or V1 *vs.* V1+PB, *p* = 0.0766; V1 *vs.* V1+PD+PB, *p* = 0.1104; Figure 3a). These results suggest differential adhesion or matrix production in mixed cultures. Mono- and co-culture biofilms of variants V2, V3, and V6 reproduced the observed results for the ancestral and V1 variant strains, indicating that the origin of the variants did not influence biofilm formation (Supplementary Figure 2a).

**Figure 3.**
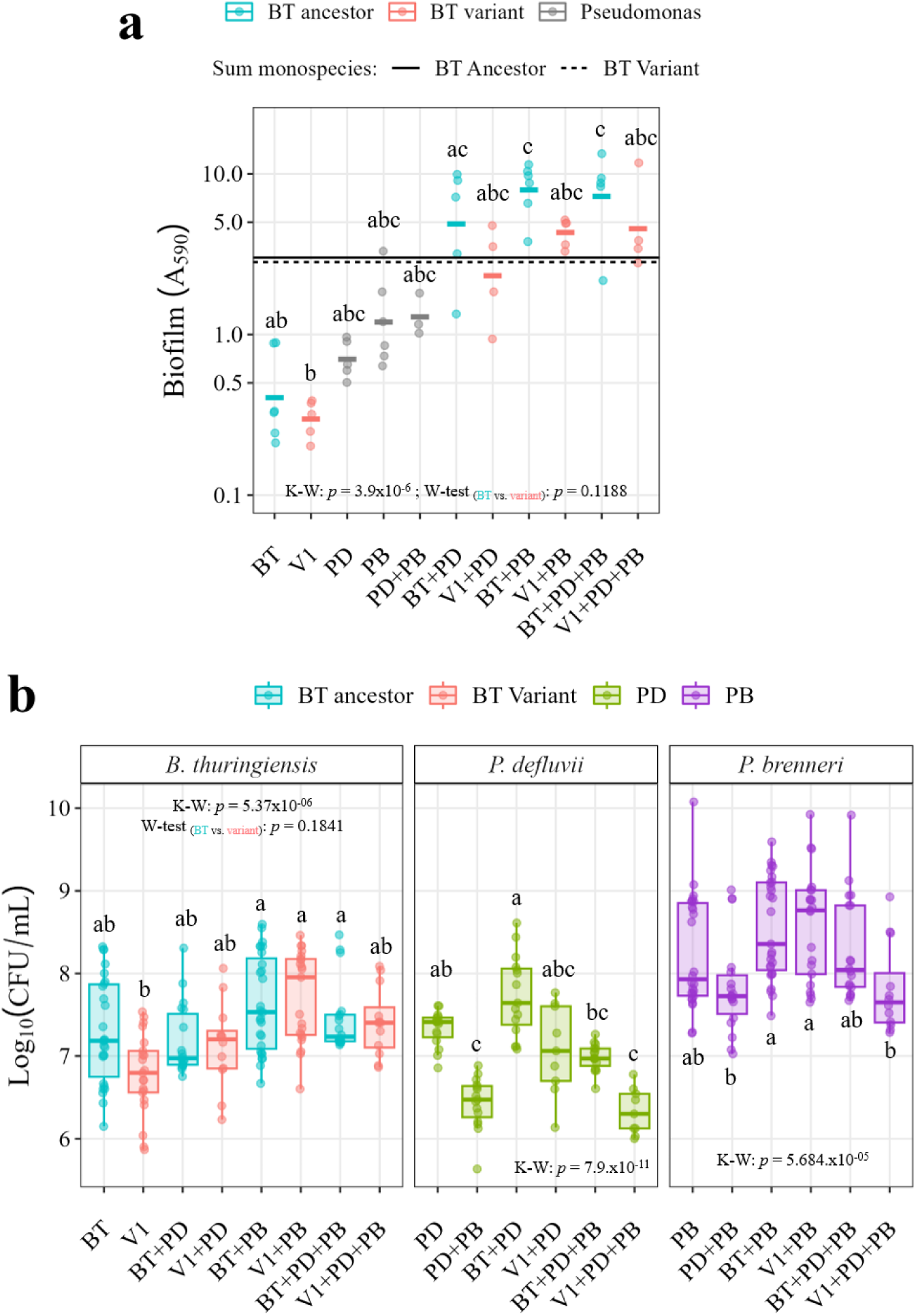
Biofilm formation of *B. thuringiensis* ancestor, V1 variant and *Pseudomonas* in mono- or mixed-species cultures after 24 h. BT: *B. thuringiensis* ancestral strain (blue); V1: *B. thuringiensis* V1 variant (red); PD: *P.defluvii*; PB: *P.brenneri*. Adhesion was measured in mono-culture biofilms or in co-culture with *P. defluvii*, *P. brenneri* or both (multispecies) in TSB medium after 24 h. K-W: significance of a Kruskal-Wallis test, testing all combinations within each panel (*p* < 0.05). Dissimilar letters are indicative of significant differences per panel, with *p* < 0.05 (Dunńs post-hoc test). W-test_(BT vs. variant)_: indicates significance of a Wilcoxońs rank sum test, comparing biomass (A_590_) or CFUs based on BT phenotype. **(a)** Mean biofilm measured as crystal violet stained biomass (A_590_). Data based on ≥ 4 biological replicates, represented as individual points. Each biological replicate consisted of 6 technical replicates. Horizontal lines represent the predicted multispecies biofilm biomass, expected in case no biofilm induction or reduction occur in co-cultures. This “predicted” value is derived from the sum of the monospecies biofilm biomass by the constituent strains. **(b)** Biofilm CFU counts per mL of *B. thuringiensis, P. defluvii* and *P. brenneri*. Data based on ≥ 9 biological replicates, represented as biological replicates, each consisting of ≥ 3 technical replicates. Dissimilar letters show significance per species.

The CV results suggested differential adhesion or matrix production in mixed cultures with the variant strain. Thus, we next quantified CFU counts for each strain in the different combinations to measure specific adhesion. Cell counts of the BT ancestral and V1 variant strains within culture condition (i.e., monoculture, co-culture, or multispecies) were not significantly different (Dunńs post-hoc test, Figure 3b, left panel), indicating similar surface colonization for both strains. However, V1 cell numbers significantly increased in co-culture with PB compared to mono-culture (V1 *vs.* V1+PB) (Dunńs post-hoc test *p* = 2.06×10^-05^, Figure 3b). Given that PB counts were unchanged despite the BT genotype, we hypothesize that the adapted variant benefitted from co-cultivation with PB. CFU counts of the other variants replicated the observations found for V1 in mono- and co-culture with PB (Supplementary Figure 2b).

In contrast, neither *Pseudomonas* species grew to a higher density in co-culture compared to mono-culture conditions. Rather, PD experienced a reduction in cell counts when co-cultured with PB or in multispecies biofilms with the variant phenotype, indicating an antagonistic effect of the adapted phenotype, and PB, on PD cell numbers (Figure 3b).

To evaluate fitness of the ancestor and the V1 variant, we performed competition experiments that included both strains, in the absence or presence of *Pseudomonas* spp. and in biofilm *vs.* planktonic settings. We mimicked the selection pressure applied in a whole serial transfer during the adaptive evolution, including co-cultivation, washing and regrowth in new media. The variant was significantly more abundant under all conditions tested, except in biofilms in the absence of *Pseudomonas* (*p* = 0.064, Dunn’s post-hoc test; Figure 4). Interestingly, under biofilm settings, the ratio of variant to wild-type phenotype increased from 4.8-fold to 25-48-fold when *Pseudomonas* spp. was present, highlighting the superior fitness of the variant in the presence of other species (Figure 4, left panel).

**Figure 4.**
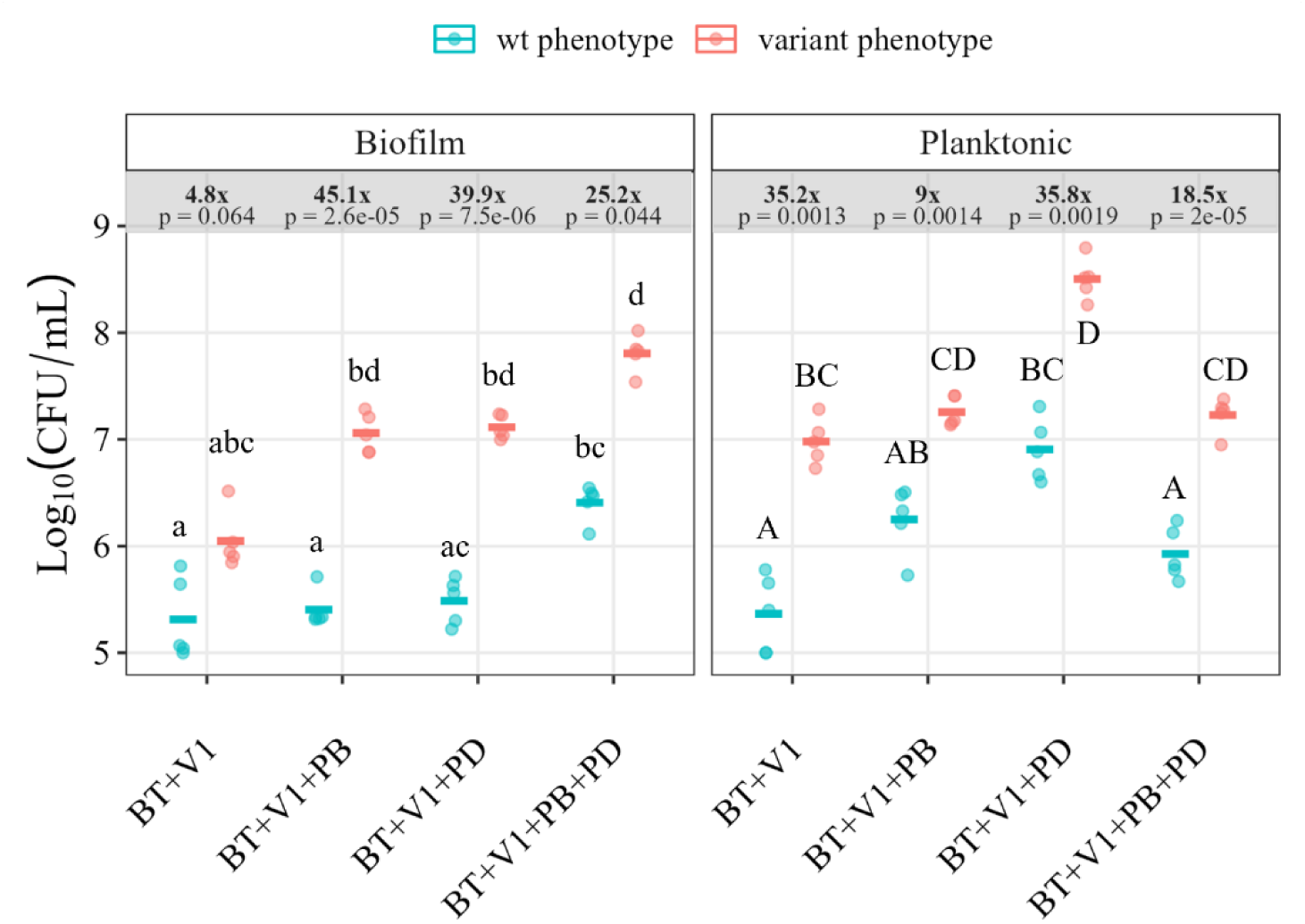
Competition of *B. thuringiensis* ancestor and V1 variant under biofilms or planktonic settings, with or without *Pseudomonas* spp. BT: *B. thuringiensis*; V1: BT V1 variant; PB: *P. brenneri*; PD: *P. defluvii*. Planktonic samples were cultivated for 24 h in TSB, diluted 1:1000 into fresh medium, and incubated for another 24 h before plating. Biofilm samples were cultured for 24 h, washed thrice, transferred to fresh media, and regrown for 24 h before plating. Plots show mean CFU counts (horizontal crossbars) of wild-type and variant colony phenotypes (Log_10_(CFU/mL)). Data represent five biological replicates (points), each with three technical replicates. Dissimilar letters denote significant difference in multiple comparisons per combination (*p* < 0.05, Dunn’s post-hoc test). Ratios of variant *vs.* wild-type phenotypes and their corresponding p-values (Dunn’s post-hoc test) are shown in grey boxes for each sample type (culture condition) and setting (biofilm or planktonic).

Although the dynamics and ratios of wild-type and variant phenotypes varied in planktonic settings, prevalence of the variant phenotype was consistently enhanced in each mixture (Figure 4, right panel). These results confirm that the variant phenotype is fitter in the short term, not just after eight days of evolution, suggesting that it outcompetes the wild-type phenotype even early in the selection process. Furthermore, we evaluated the impact of the specific *Pseudomonas* spp. on the selection of variant or wild-type phenotypes in a triple culture system under both planktonic and biofilm settings. The abundance of the wild-type phenotype was significantly lower in biofilm settings compared to planktonic in the presence of PB, whereas abundance of the variant phenotype was unchanged (Two-sample t-test: *p*wt = 0.0119, *p*variant = 0.0923; Supplementary Figure 3a). In contrast, the abundance of both the wild-type and variant phenotypes was enhanced in planktonic conditions compared to biofilms when PD was present (Two-sample t-test: *p*wt =2.17×10^-05^, *p*variant = 0.00794; Supplementary Figure S3b). These results suggest that PD and PB interact differently with *B. thuringiensis*, which may result in different selective pressures posed by the two *Pseudomonas* spp. Specifically, PD may facilitate nutrient availability or alter competition dynamics in ways that favor planktonic growth of *B. thuringiensis*, regardless of its phenotype.

In conclusion, these findings suggest that the evolutionary adaptations in these variants may specifically affect biofilm matrix production and aggregation without significantly impairing their ability to adhere to surfaces. The observed reduction in matrix production in mixed-species biofilms with the variant could stem either from the inability of *B. thuringiensis* light variant to induce matrix production in *Pseudomonas* spp. or from intrinsic diminished matrix production in such variants.

### Matrix glycan composition altered by co-cultivation and BT light variant

We observed a synergistic effect on biofilm biomass in mixed species, compared to mono-culture, though only significantly different with BT ancestor (Figure 4a). However, no correlation was found between such induction and CFU counts of either of the strains. We hypothesized that such differences in biofilm biomass between ancestral and variant strains in mixed species might imply altered production of biofilm matrix, either in BT variants or in the *Pseudomonas* species. Thus, we investigated the influence of interspecies interactions and/or acquired mutations on the amount and types of glycans within the biofilm matrix.

We screened multispecies biofilms (BT+PD+PB) with 78 different fluorescent lectins binding to carbohydrates of varying nature (Supplementary Table 5). After filtering the binding results by fluorescence signal (data not shown), three lectins with strong binding in multispecies biofilms were identified for further analysis: AAL, WGA, and RCA. AAL binds α-fucose-containing polymers, WGA binds N-acetylglucosamine (GlcNAc), and RCA binds galactose and N-acetylgalactosamine (Gal/GalNAc) glycans [45]. Considering the minimal difference in biofilm biomass between co-culture and multispecies biofilms (Figure 3a), we used co-culture conditions as it closely resembled multispecies conditions. Therefore, we assessed binding of the three lectins to BT mono- and co-culture biofilms with PB, for both ancestral and variant strains.

AAL lectin bound to BT cells and certain matrix-like structures in BT mono-culture biofilms, while the variants displayed reduced lectin signal (Figure 5a). This indicates that the composition of glycoconjugates in the cell wall or surface differed between the variants, leading to differential AAL binding. Image quantification of lectin binding supported visual observations, significantly reducing lectin binding in the variants (Figure 5b). However, the amount of lectin binding per biofilm volume (biovolume) unit was not significantly changed in the variants (Figure 5c). We cannot exclude the possibility that intrinsic heterogeneity of biofilm and differences among slide replicates, even within biological replicate, could account for these results. Moreover, some of the replicates also showed some unspecific lectin binding that could influence the analysis.

**Figure 5.**
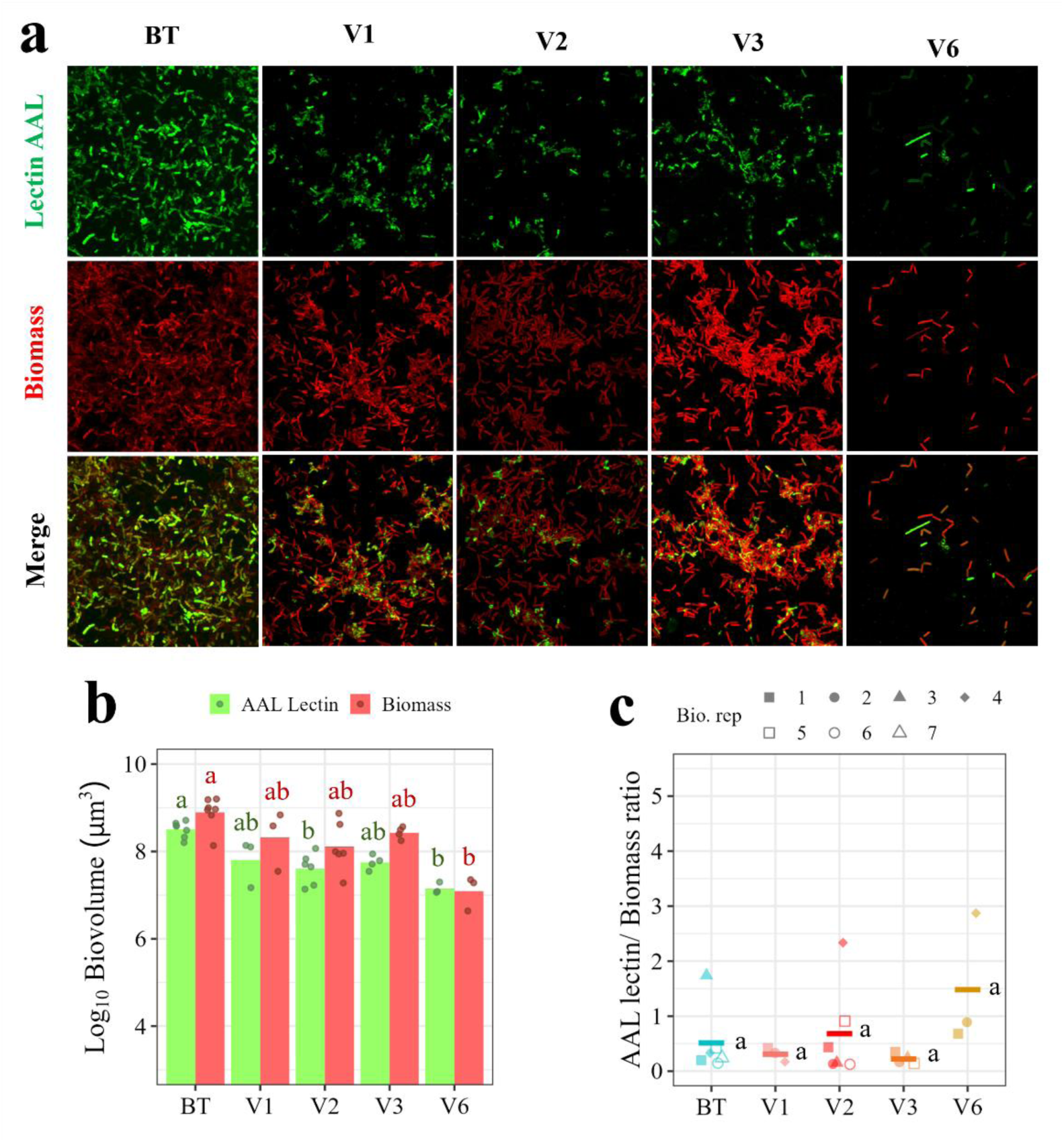
AAL lectin staining in mono-culture biofilms of *B. thuringiensis* ancestor and variant strains. BT: Ancestral *B. thuringiensis* strain; V1: variant1, V2: variant 2; V3: variant 3; V4: variant 4; PB: *P. brenneri*. Biomass: red fluorescent signal of biofilms stained with the cell biomass stain SYTO60. Lectin AAL: green fluorescent signal of biofilms stained with the fluorescent lectin AAL-FITC. Merge: combined fluorescent signal of cell biomass and lectin stains. **(a)** Maximum intensity projection (MIP) images of Z-stacks recorded by confocal laser scanning microscopy for 24-hour biofilms. The scale of the images is 124 µm across. **(b)** Biofilm volume (Biovolume, μm^3^) of lectin (green) and cell biomass fluorescence (red). Biovolume was calculated using BiofilmQ [86] per channel, based on ≥ three biological replicates (represented as points) and two images per replicate. Bars show mean values per channel. Dissimilar letters denote significant p-value (Dunńs post-hoc test, *p* < 0.05) per channel (green = lectin; red = cell biomass). **(c)** Ratios of lectin biovolume/ biomass biovolume for each strain. Data based on ≥ three biological replicates, represented by different symbols, and two images per replicate. Crossbars indicate the mean ratio for each strain. Dissimilar letters denote significant p-value (Dunńs post-hoc test*, p* < 0.05).

Co-cultivation of BT ancestral or variant strains with PB however showed similar AAL binding, (Supplementary Figure 4), suggesting a change in glycoconjugates triggered by interspecies interactions. Moreover, PB also showed AAL binding in mono-culture and, thus, association of lectin signal with the specific matrix producer may be cumbersome in co-culture conditions (Supplementary Figure 4, BT and PB panels).

Unlike AAL lectin binding, no differential binding was observed for RCA lectin, regardless of the BT genotype or cultivation (mono- *vs.* co-culture with PB). Moreover, in PB biofilms, this lectin bound thin filaments that revealed a galactose/N-Acetylgalactosamine polymer network (Supplementary Figure 5a). On the other hand, WGA bound to BT cell surfaces (Supplementary Figure 5b), likely the cell wall because this lectin binds to the sugar GlcNAc, which is a major component of peptidoglycan [46]. However, there was no visible difference observed in either the variants or mixed-species combinations (data not shown).

In conclusion, AAL lectin staining showed a tendency of reduced production of α-fucose polymers by BT variants but lectin-biomass ratios were not conclusive. However, other matrix components may be influenced by the distinct phenotypic biofilm formation by BT genotype, such as protein abundance.

### Matrix protein abundance of BT light variant decreased in response to co-cultivation and genotype

We next investigated the effects of interspecies interactions and acquired mutations in BT on the proteomes of biofilm matrices, hoping to explain the reduced biofilm matrix production observed in either the BT variants or the *Pseudomonas* species. We specifically examined variants V1 and V6 due to their distinct origin of isolation (V1 from co-culture with PB and V6 from mono-culture), to eliminate the possibility that the isolation origin influenced the abundance of matrix proteins. Our proteomic experiments utilized a chemical extraction of biofilm matrix as input material, which results in a lower number of expected proteins compared to total biofilm samples.

First, we performed a comparative analysis of BT proteomes by genotype (variants *vs*. ancestral strains) and culture condition (mono-culture *vs.* co-culture with PB) (Figure 6). It is important to note that co-culture proteomes likely contain BT and PB derived proteins from the extracted matrices. The analysis showed that BT ancestor mono-culture samples were the most distinct from the rest, clustering together on the right of PC1 (Figure 6a), as evidenced by probabilistic principal component analysis (PPCA). The variation in PC1 was driven by the BT genotype (ancestor *vs.* variant), while the cultivation condition (mono-culture *vs.* co-culture) determined PC2. Light variant co-culture or mono-culture samples could not be discriminated, while BT ancestor co-culture samples were grouped in a distinct cluster compared to mono-culture. Interestingly, BT co-culture proteomes overlapped with variant mono- and co-culture proteomes. Consistently, comparison of BT ancestor *vs.* variants in mono-culture evidenced the largest difference in proteomes. Only 20 proteins out of 187 were common to all three BT strains (Figure 6c and Supplementary Table 6) and 11 significantly changed, related to amino acid and carbohydrate metabolism energy production, among others. In contrast, 150 proteins were exclusively detected in the ancestor (Supplementary Table 7), indicating a major reduction in protein abundance in both variants. Among those, different proteins involved in sporulation or regulated by the Stage 0 sporulation protein, Spo0A, were only detected in the ancestor, consistent with sporulation-defective phenotypes observed in the variants.

**Figure 6.**
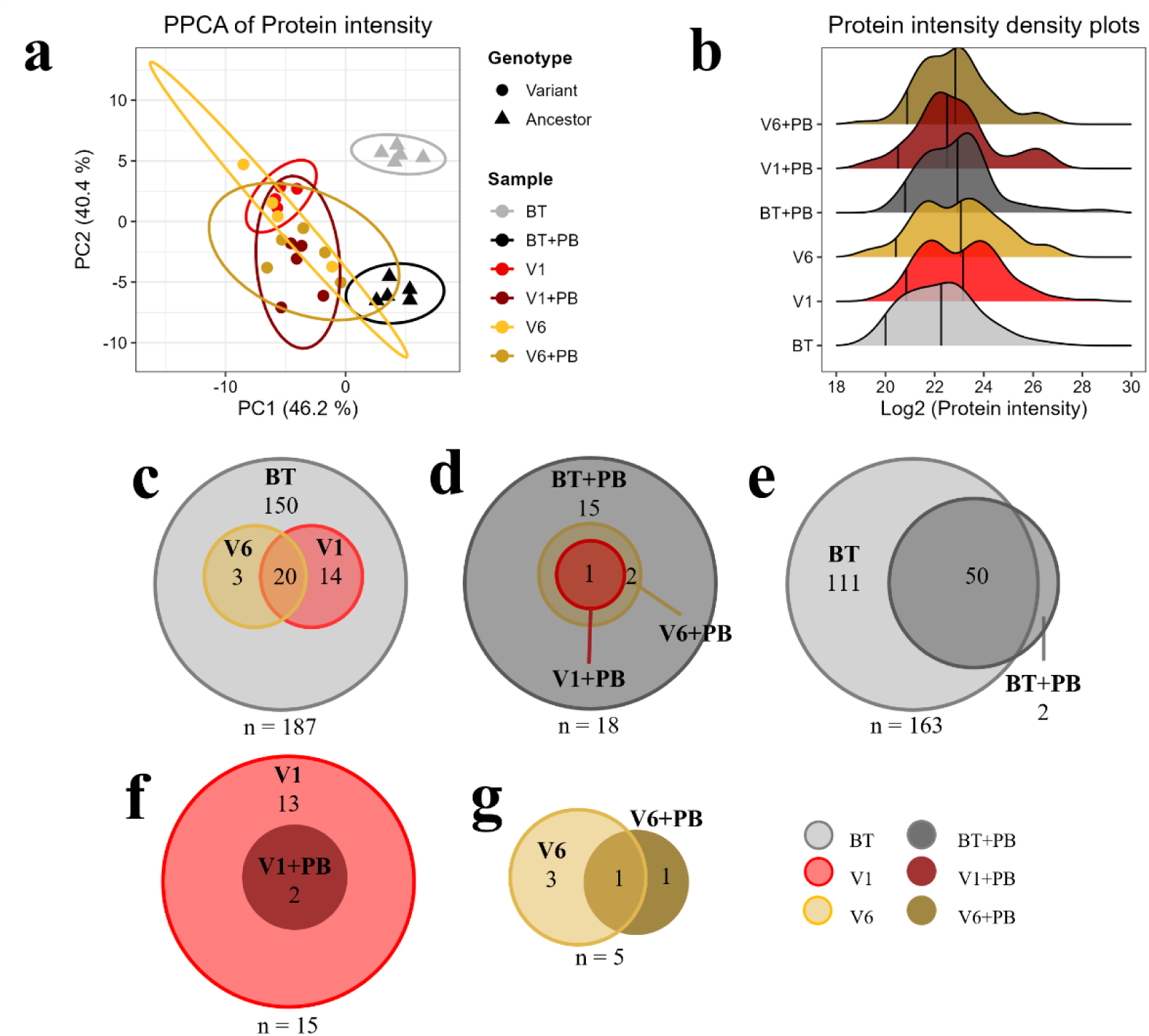
Differential protein abundance in *B. thuringiensis* ancestor *or variants* in mono-culture *vs.* co-culture biofilms with *P. brenneri*. BT = Ancestral *B. thuringiensis* strain; PB = *P. brenneri*; V1 = *B. thuringiensis* variant 1; V6 = *B. thuringiensis* variant 6. **(a)** Probabilistic PCA plot of all monoculture and co-culture proteomes, based on Log_2_ transformed protein intensity data with 5 biological replicates per sample. Color indicates the sample type (strain and culture condition) and shape the genotype: ancestral (triangles) or variant (circles). Ellipses indicate sample group clustering based on the principal components. **(b)** Density plots of Log_2_ transformed protein intensity for mono-culture *vs.* co-culture samples representing 5 biological replicates. Percentile 0.05 is indicated as the leftmost vertical line, while middle lines denote protein intensity means for each sample excluding NA values (no protein detected). (**c-g)** Venn diagrams of *B. thuringiensis* mono-culture proteomes (c), co-culture proteomes (d), or mono-culture *vs.* co-culture proteomes (e-f), considering proteins detected in ≥ 60 % of each sample (3/5 biological replicates). **(c)** Mono-culture proteomes **(d)** Co-culture proteomes **(e)** BT *vs.* BT+PB proteomes **(f)** V1 *vs.* V1+PB proteomes **(g)** V6 *vs.* V6+PB proteomes. n = total proteins detected per comparison.

To discard unique expressions due to detection limits, we calculated the mean and 0.05 percentile of each proteome to use as a cutoff (Figure 6b and Supplementary Table 8). However, DivIVA, SpoVG, immune inhibitor A (CDS_04360 and CDS_05099), neutral protease B (CDS_00401) and the major biofilm matrix component TasA(CDS_04363 and CDS_04365) were still above the expression cutoff in the ancestral BT strain, compared to the light variants. Remarkably, the four latter proteins were also more abundant in the ancestral co-culture (Supplementary Table 9), suggesting that their abundance is influenced by BT genotype —ancestor — and the presence of PB.

Co-cultivation of PB with BT ancestral or variant strains reduced biofilm matrix protein abundance by 90 % (17 proteins co-culture *vs.* 187 in mono-culture), suggesting that PB minimized the differences of BT proteomes (Figure 6d). However, 15 proteins were uniquely detected in the ancestor co-culture, including the four mentioned above (Supplementary Table 9). Mono-culture *vs.* co-culture comparisons rendered highly different results depending on the genotype compared. In the ancestor pairwise analysis, 111 out of 163 proteins were only detected in mono-culture (Figure 6e and Supplementary Table 10), supporting the previous observation that PB decreased abundance of biofilm matrix-associated proteins in BT. However, comparisons between the V1 and V6 variants resulted in reduced matrix proteomes, with 15 and 5 proteins detected, respectively (Figures 6f-g and Supplementary Tables 11-12). Importantly, these findings reflect the proteomes of extracted matrices rather than global proteomes, as we would anticipate a broader range of differentially abundant proteins in global proteomes due to *spo0A* mutations, considering its pivotal regulatory role.

We hypothesize that the acquired mutations may switch off the expression of multiple genes involved in sporulation, a highly complex and tiered-regulated process linked to biofilm formation. There is previous evidence that *spo0A* mutations in *Bacillus subtilis* enhanced maintenance metabolism [47] and increased glucose consumption and acetate formation [48]. This would be consistent with the faster doubling time observed for the variants, given that fewer proteins and consequently a lower amount of energy are required in the absence of sporulation.

We investigated whether matrix proteins were differentially abundant in PB based on culture condition and BT genotype (Supplementary Figure 6). Out of 169 detected proteins, mono- and co-culture proteomes showed similar mean abundance and 0.05 quantiles (Supplementary Table 8 and Supplementary Figure 6a). Unlike BT results, co-cultivation did not strongly affect the number of detected proteins, with 64% shared among all samples (n = 108) (Supplementary Table 13 and Supplementary Figure 6b). PB mono-culture samples clustered separately to co-culture samples, indicating that cultivation drove PC1 (Supplementary Figure 6c). Seventeen proteins were uniquely detected in co-culture with BT ancestor (Supplementary Table 13), though co-culture samples of BT, V1, or V6 could not be discriminated in the PPCA. While no unique or differentially abundant matrix proteins were found in PB, multiple proteins involved in amino acid transport and metabolism varied based on culture condition. Co-cultivation reduced the abundance of several amino acid transporters in PB regardless of BT genotype (arginine, branched-chain, cysteine, glutamate/aspartate, and putrescine, Supplementary Figure 6d). Other proteins involved in branched-changed transport (braC_1), glycine, tyrosine or arginine/ornithine metabolism were only detected in PB monoculture (Supplementary Table 13). These findings suggest that PB optimizes amino acid utilization differently depending on environmental conditions. In co-culture with BT, ecological dynamics might shift, altering metabolic priorities and protein abundance. This aligns with our previous observation that V1 variant CFU counts increased in co-culture with PB, suggesting that the variant benefits more from co-cultivation with PB than the ancestral strain (Figure 4b).

Based on all proteomics results, we hypothesize that the lower abundance of matrix-associated proteins in BT light variants correlates with the previously observed reduction in biofilm biomass in the variant mixtures. Furthermore, the absence of differential or unique abundance of matrix proteins in PB suggests a deficiency in BT’s production of matrix components associated with *spo0A* mutations. These results are also consistent with the observed tendency of reduced AAL lectin binding in the variants.

## Discussion

Microbial interactions influence the fitness, survival, and evolution of interacting species, enabling better resource utilization, mitigating competition, enhancing mutual dependencies, or optimizing nutrient exchange. Despite their predominance and significance, adaptive evolution experiments focusing on multispecies biofilms remain limited. This study aimed to address this gap by examining effects of interspecies dynamics and spatial structuring on trait selection within the Gram-positive bacterium BT.

Here, we used short-term evolution assays where BT, PD, and PB were cultured individually or in combination, under both planktonic and biofilm conditions (Figure 1a). Throughout, these assays, a distinct phenotypic variant of BT, which we termed “light variant”, consistently emerged. Notably, this variant was more prevalent than the wild-type phenotype in biofilm environments (Figure 1b), and its selection further amplified by interactions with *Pseudomonas* species—indicating a potentially fitter phenotype under such conditions. While previous experimental evolution studies in biofilms have documented the emergence of adapted variants exhibiting enhanced biofilm characteristics relative to their ancestors, our findings diverge [18, 20, 49, 50]. For instance, Lin and collaborators identified a “fuzzy spreader” phenotype of *B. thuringiensis* in a bead-transfer evolution study, where all adapted lineages resulted in enhanced biofilm production compared to its ancestor [26]. In contrast, the BT variants identified in our study produced less biofilm within mixed-species contexts, which was associated with a reduction in biofilm matrix components like TasA. Despite this, the adapted phenotype prevailed across different environments (planktonic or biofilm, Fig. 1b), suggesting a fitness advantage over the ancestral form. Moreover, the adapted variant displayed reduced auto-aggregation compared to the ancestral strain (Figure 2a-b). This decrease in aggregation could optimize the distribution of cells within the environment, facilitating better access to nutrients and, possibly, enhancing survival and growth. This advantage is consistent with the observed shorter generation time of the non-sporulating variants and higher prevalence of the light variant V1 when co-cultured with the ancestor, especially in the presence of other species (Figure 4). These traits together would confer a substantial fitness benefit, especially in competitive environments with other species, where the ability to outcompete other organisms and rapidly access resources could be crucial for survival and proliferation (Figure 7). However, considering the comparable biofilm characteristics between the ancestor and the variants, our findings suggest that BT adaptation in biofilms may be driven by factors beyond adhesion or biofilm formation, such as nutrient competition and subsequent metabolic optimization, which may also be more compatible with *Pseudomonas* co-existence.

**Figure 7.**
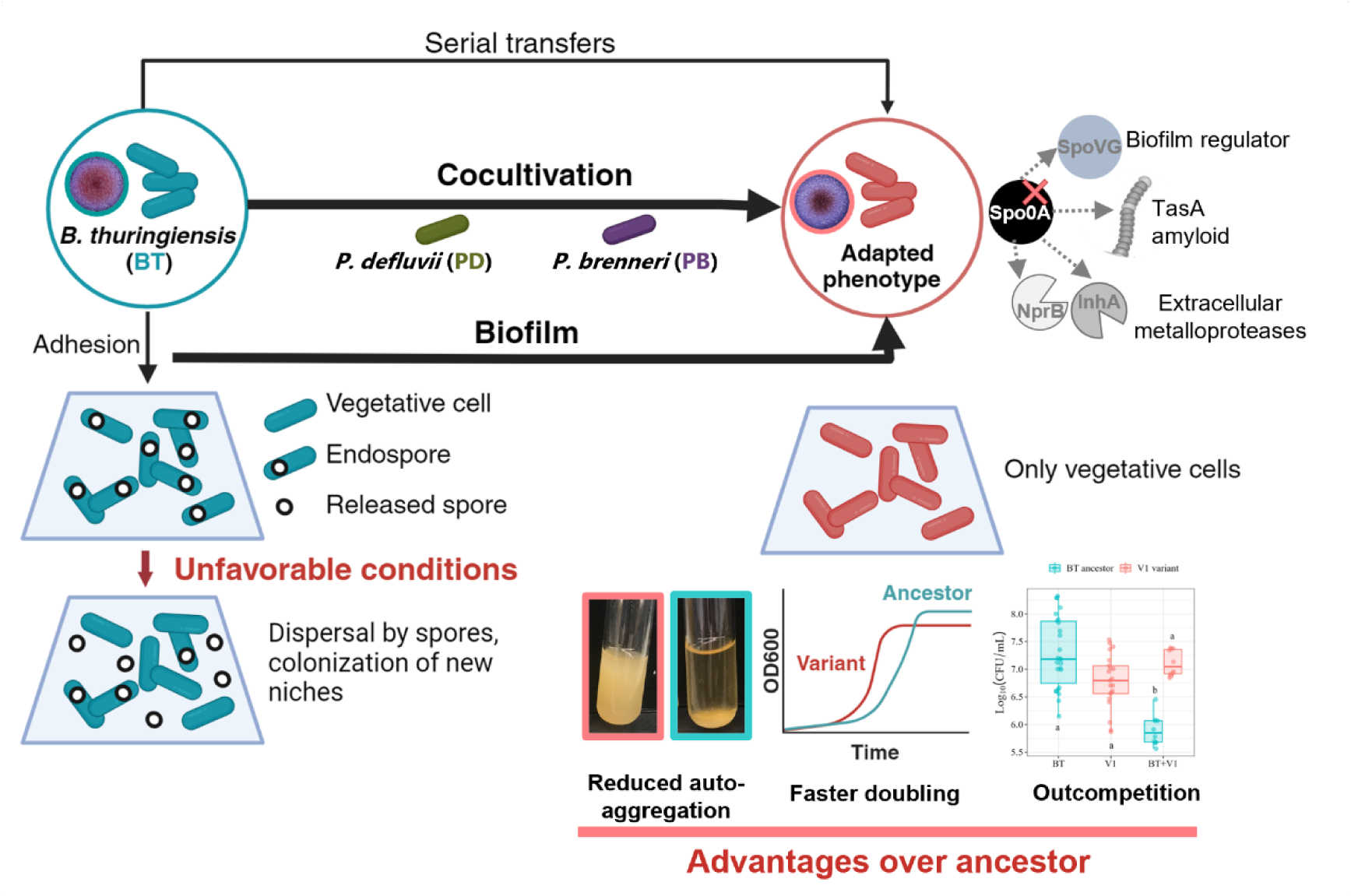
Proposed model for adaptation of the evolved variants. *B. thuringiensis* non-sporulating variants emerged in all conditions, but biofilm settings and co-cultivation with either or both *P. brenneri* and *P. defluvii* enhanced selection of the variant (thicker arrows). In adverse conditions, spores might aid in dispersing biofilms in the ancestral strain. However, adapted variants, lacking sporulation, could experience restricted dispersal. The adapted variant lacks sporulation, displays reduced auto-aggregation, faster doubling time than its ancestor and can outcompete it in co-culture biofilms. Variants lack proteins SpoVG, the matrix amyloid TasA and two extracellular metalloproteases involved in matrix recycling. Indirect Spo0A regulation is indicated by dashed arrows [41, 54, 67, 68, 73, 99].

Biofilm formation and spore production are two different survival strategies of bacteria facing adverse environmental stresses. However, intertwined regulatory pathways between biofilm formation and sporulation have been proposed for both *B. subtilis* and *Bacillus cereus* [51, 52]. *spo0A* mutants in other *Bacilli* such as *Bacillus velezensis*, *B. subtilis* or *B. cereus* are impaired in biofilm formation, since Spo0A de-represses EPS production and promotes biofilm [53–56]. Whole-genome sequencing of the variants linked lack of sporulation with mutations in key genes regulating sporulation (Supplementary Figure 1). V1, V3 and V6 variants harbored mutations in *spo0A*, which has a central role in sporulation [41, 42]. However, BT variants lacking Spo0A showed comparable CFU counts and matrix production to the ancestor in mono-culture (Figure 3). Our results demonstrate that, even though the variant displayed reduced capacity for auto-aggregation (surface-independent aggregation), it does not entirely lose the ability to attach to surfaces, distinguishing it from other *spo0A* mutants in Bacilli such as *B. subtilis* and *B. cereus*. On the other hand, V2 variant showed structural variation (SV) affecting the *soj* gene. SV can confer change in gene copy number, creation of new genes, altered gene expression and many other functional consequences [57]. Other studies have demonstrated that certain mutations creating structural variants can enable new adaptive phenotypes to arise in laboratory populations of microorganisms, that may be inaccessible single-nucleotide mutations [43, 58]. Soj has been associated to correct chromosome segregation and partitioning during both growth and initiation of sporulation in *B. subtilis* [44]. The regulation of chromosome structure by Soj is particularly important during sporulation, when the bacterial cell undergoes significant changes in chromosome architecture. Thus, structural variation affecting this gene may explain the non-sporulating phenotype of this variant.

A recent study established a link between spore formation and biofilm dispersal in *B. subtilis* [59]. Under unfavorable conditions, spores may detach from the biofilm, dispersing through air, water, or other means, and potentially establishing new biofilms elsewhere, thereby promoting bacterial survival and spread. Our findings indicate that, under the controlled conditions of the laboratory where environmental stressors are minimal or absent, sporulation might become dispensable for BT (Figure 7). However, non-sporulating variants have also been reported in other Bacilli in more relevant ecological contexts, such as deep nutrient or energy deprivation, food surfaces or insect cadavers, where they survived from days to months [60–62]. This suggests that our findings can have ecological relevance and that such mutations can be selected and fit in more extreme conditions.

Irrespective of the specific mutations, all BT light variants shared identical phenotypic traits: grew faster than the ancestor (Supplementary Table 3), displayed reduced auto-aggregation (Figure 2a-b), did not sporulate (Figure 2c), and bound the AAL lectin less, which pointed towards decreased matrix glycoconjugates (Figure 5). Moreover, proteomic analysis revealed that different matrix proteins were uniquely detected in the ancestor. Among these, two TasA/CalY orthologues and SpoVG were detected in BT ancestor but not in the variants (Supplementary Table 7). TasA is a major matrix component in *Bacillus* biofilms and its absence results in significant defects in biofilm and pellicle formation in *B. subtilis* [63, 64]. Its paralog in *B. cereus* is needed for root colonization, while in *B. thuringiensis*, the TasA-orthologue CalY has been identified as a cell-surface adhesin and described as a major virulence factor and biofilm matrix protein [65, 66]. Interestingly, both CalY orthologues (*tasA_3* and *tasA_4*) are located in the same operon as *sipW*, regulated by SinR and, consequently, indirectly controlled by Spo0A in *B. cereus* (Supplementary Figure 7). Such regulatory circuit would explain the lack of TasA in *spo0A*-deficient variants. On the other hand, SpoVG plays a critical role in the spore-coat formation of *Bacillus anthracis* and is a regulator of both sporulation and biofilm formation in *B. cereus* 0-9 that activates the levels of Spo0A and, thus, the fate of *B. cereus* cells: biofilm or sporulation [67–69]. Proteomic results also evidenced the unique detection of two metalloproteases only in BT ancestor: NprB and InhA (immune inhibitor A), linked with protein recycling during sporulation and indirectly regulated by Spo0A [41, 70, 71]. InhA coding gene, *ina_4*, is also located in the same genomic region than the two *calY* orthologues and regulated by the SinI-R regulon in other Bacilli [72, 73]. We speculate that the lack of expression of TasA and the biofilm regulatory protein SpoVG could contribute to the differences observed between CFU counts and total biofilm biomass for the variant phenotype (Crystal Violet).

Apart from BT genotype, co-cultivation was another factor influencing protein abundance, yet differently depending on the target species. Co-cultivation significantly altered the proteome of BT but only in the ancestral strain (Figure 6a). Meanwhile, for PB the presence of BT greatly impacted protein abundance regardless of the phenotype (Supplementary Figure 6c), showing consistent lower abundance of amino acid transporters in PB co-culture (Supplementary Figure 6d). This could indicate competition for resources or utilization or distinct metabolic pathways in co-culture. We have previously reported that cooperative interactions between community members can facilitate metabolic cross-feeding on specific amino acids, though in a different multispecies biofilm model [74, 75]. Additionally, previous studies have shown that coexistence can remain stable over time, for instance, by oppositely altering the pH or secreting costless by-products to the other species’ benefit [18, 76–80]. Most reported *Bacillus-Pseudomonas* co-cultures demonstrate either competition or amensalism, driven by bioactive natural products, while positive interactions are usually associated with cross-feeding or shared resources [31]. Our evolution experiment revealed significant reduction of PB CFU counts in planktonic co-culture with BT, unlike biofilm conditions, suggesting that spatial structure facilitates co-existence of these species. Furthermore, higher abundance of the variant phenotype was observed in co-culture with PB in both planktonic and biofilm settings, suggesting that the adapted variant benefits from PB, compared to its ancestor. This pattern is consistent with a non-exploitable dynamic, with the BT variant benefiting through improved growth without negatively impacting PB.

In summary, our study evidences the ecological role of non-sporulating variants of *B. thuringiensis* in multispecies communities, demonstrating that prevalence of the variant phenotype is likely influenced by its lower auto-aggregation capacity, its competitive advantage in the presence of *Pseudomonas* spp. and its ability to thrive in both planktonic and biofilm environments (Figure 7). However, our results showed enhanced prevalence of the variant phenotype over the wild type in biofilm settings and in the presence of *Pseudomonas* species, suggesting that selection of the non-sporulating variants is strongly influenced by such factors (Figure 7). Our findings challenge the traditional view that biofilm evolution invariably selects for stronger biofilm-forming phenotypes. Instead, we demonstrate that adaptation may favor traits that optimize resource competition or metabolic efficiency over biofilm reinforcement. These results have important ecological implications, as similar evolutionary dynamics may occur in natural microbial communities where competition and cooperation among species influence community structure. Hence, this study evidences the importance of considering population diversity—both intra- and interspecific, along with spatial structure as key determinants of bacterial adaptation.

## Methods

### Strains and materials

The three bacterial strains PD, PB, BT were isolated from water samples collected at a wastewater treatment plant in Kalundborg, Denmark, and their genome s were sequenced in another study [29]. All strains were grown at 24 °C in Tryptic Soy Broth (TSB; 17 g pancreatic digest of casein, 3 g papaic digest of soybean meal, 5 g sodium chloride, 2.5 g dextrose, 2.5 g dibasic potassium phosphate in 1 L distilled water, pH 7.3), supplemented with 1.5 % agar when needed. To distinguish the bacterial strains and/or species variants, 40 µg/mL of Congo Red (Merck) and 20 µg/mL Coomassie Brilliant Blue G (Sigma-Aldrich) were added to the TSB base, referred to as TSB Congo Red plates. In previous studies, this media has facilitated identification of adapted variants and differentiation among species [18, 77].

### Short-term evolution assay with spatial structure

Overnight cultures in TSB medium were adjusted to an optical density at 600 nm (OD_600_) of 0.15 in fresh TSB prior to usage. The experiment was conducted in 24-well plates, where each well contained a polycarbonate chip (12 x 12 x 0.78 mm^3^) (PC) that was diagonally tilted so that bacteria could adhere to both sides of the chip. In each well, 2 mL of OD_600_ 0.15 cultures were added, either as mono-species culture or mixed-species culture. For mixed-species conditions, 1:1 or 1:1:1 OD_600_ ratios of the species were used, that is, 1 mL or 0.66 mL of each species at OD = 0.15, for dual- and multispecies, respectively. Every 24 hours (h), the floating pellicle was gently removed and discarded before the polycarbonate chip, containing adhered cells, was transferred to a new well, and the chip was washed thrice in 1x phosphate-buffered saline (PBS) to remove any leftover planktonic/pellicle cells. This was done consecutively for 8 days, followed by washing the PC chips before transferring to a 5-mL Eppendorf tube containing 2 mL 1x PBS. The biofilm was then removed from the PC surface by shaking at 600 rpm for 5 minutes (min) (Titramax1000, Heidolph). The resulting suspension was then plated on TSB Congo Red plates with three technical replicates to estimate colony forming units (CFUs) and to screen for new variants with different colony morphology.

### Short-term evolution assay without spatial structure

To test selection pressure in conditions not favoring spatial structure, a short-term evolution assay was also performed with well-mixed planktonic cultures. Cultures were adjusted to OD_600_ = 0.15 as described above, and 5 mL of this culture was incubated overnight, with shaking at 250 rpm (KS 250 basic, IKA). After 24 h, 5 μL of this culture was transferred to new fresh media. The transfer was repeated for 8 days, followed by plating on TSB Congo Red plates with three technical replicates.

### Sedimentation dynamics in planktonic cultures of BT ancestor and V1 variant

The sedimentation dynamics of the BT ancestor and V1 variant strains were assessed in planktonic cultures grown in TSB at 24 °C and 250 rpm for 20 h. A 1-mL aliquot of each strain culture was transferred to a spectrophotometer cuvette. Culture aliquots were maintained under static conditions to observe natural sedimentation patterns. OD_600_ was measured at time zero (t0) and for 60 minutes (min) in 5 min intervals using a GENESYS UV10 spectrophotometer (Thermo Scientific), to quantify sedimentation. Values were expressed as the percentage of initial OD_600_ (t0). Measurements were performed for three biological replicates of each strain, and mean percentage of initial OD_600_ calculated per time point.

### Microscopic analysis of auto-aggregation in planktonic cultures of BT ancestor and V1 variant

The auto-aggregative properties of the BT ancestor and V1 variant strains were analyzed using fluorescence microscopy. Planktonic cultures of both strains were grown in 5 mL TSB for 20 h at 24 °C and 250 rpm. Cultures were left for 1 h without shaking and 20 µL of top or bottom phases stained with 10 µM SYTO 16 (nucleic acid stain) to visualize cellular distribution. Images of the top and bottom phases of the cultures were captured using a Zeiss Axio Observer Z1 epifluorescence microscope equipped with a LD Plan-NEOFLUAR 20x objective (numerical aperture 0.4).

### Quantification of biofilm formation

Assessment of the biofilm formation on the polycarbonate surfaces was performed using a modified crystal violet (CV) assay [81]. Biofilms were allowed to grow on PC chips for 24 h as described above, washed thrice in 1x PBS and then transferred to a well of a 24-well plate containing 2 mL of 1 % CV aqueous solution and incubated for 20 min in the dark. Then, the chips were carefully removed and rinsed five times in 1x PBS. CV-stained chips were then incubated in 2 mL of 96 % ethanol for 30 min. The biofilm biomass was then quantified as the absorbance of CV solution at 590 nm (A_590_). To avoid measurement saturation, samples were diluted in 96% ethanol when A_590_ was higher than 1.

### Growth curves in liquid cultures

Cultures of BT strains were grown overnight in liquid TSB medium. Then, 1:100 dilutions of the overnight cultures in TSB were prepared, and these cultures were used to inoculate 96-well plates, using 3 biological and 3 technical replicates. Growth was monitored for 24 h at 24 °C with 282 rpm double orbital shaking and OD_600_ readings every 10 min. Growth data was integrated to calculate different growth parameters using the Growthcurver R package [82] to determine different parameters. These included: t_mid, the point of inflection of the growth curve; K, carrying capacity or maximum size of the population; t_gen, generation time of a strain; r, intrinsic growth rate of a population; sigma, the fit goodness of the model. The resulting calculations can be found in Supplementary Table 3.

### Whole-genome sequencing and analysis of variants

BT ancestor and isolated variants V1, V2, V3, and V6 (Supplementary Table 2) were genome-sequenced by extracting the DNA using DNeasy Blood & Tissue Kit (Qiagen) from overnight cultures. The samples were sequenced with short reads using Truseq libraries on the Illumina NovaSeq 6000 and 2×150 cycles (Novogene, UK).

Data was analyzed using breseq (v0.35.0) to call variants using the –p polymorphism flag when analyzing population sequences [83]. As the reference genome, we used the previously constructed hybrid assembly of long and short reads from BT, available online (NCBI) with ID JARTVI000000000. Mutations detected between our ancestral strain and the reference genome were not considered. The breseq software also predicts new junctions within the genome which may occur due to mobile element insertions, large deletions, structural rearrangements and prophage excision. These analyses were also included. A full list of the mutations can be found in Supplementary Table 4.

### Optical microscopy

Liquid cultures of the different strains were incubated for 24 h in TSB at 24 °C while shaking at 250 rpm. From these 24-h cultures, 10 μl were pipetted onto a microscope glass slide, and a coverslip was applied on top. Samples were observed with an Olympus BH2 optical microscope with 10x and 40x magnification objectives, with 0.25 and 0.65 numerical apertures, respectively.

### Sporulation assay

To compare the sporulation capacity of *B. thuringiensis* and its variants, a sporulation assay was performed similarly to the description by Toukabri et al. [84]. Firstly, overnight cultures (24 h) of the ancestor and variants were serially diluted and plated on TSB Congo Red to quantify the concentration vegetative cells in each culture. Similarly, an aliquot of such cultures was heat-shocked at 80 °C for 20 min before plating on TSB Congo Red plates to quantify sporulating cells. Plates were incubated two days at 24 °C before counting. This was repeated for 3 consecutive days (48-, 72- and 96-h cultures) to assess whether incubation time influenced spore formation. Furthermore, the sporulation assay was performed for the biofilms grown on PC chips to estimate the number of spores produced in mono or mixed-species cultures.

### Competition assay of BT ancestor and V1 variant in biofilm and planktonic settings with or without *Pseudomonas* spp

To assess fitness of the ancestral BT and V1 variant, they were grown in the absence or presence of *Pseudomonas* spp. under planktonic or biofilm conditions simulating a full serial transfer of the evolution experiment. Overnight cultures of each strain were adjusted to OD_600_ = 0.15 in fresh TSB and mixed in equal volume ratios: 1:1 (BT+V1), 1:1:1 (BT+V1+PB; BT+V1+PD) and 1:1:1:1 (BT+V1+PB+PD).

Inoculation of biofilm and planktonic samples was done as detailed in the short-term evolution assay with and without spatial structure, respectively, however with only one daily transfer. Specifically, planktonic samples were cultivated in TSB for 24 h at 24 °C and 250 rpm, diluted 1:1000 in fresh medium and incubated for another 24 h before plating. For biofilm samples, they were cultivated in TSB for 24 h at 24 °C under static conditions, washed three times with fresh medium, transferred to new media, and regrown for another 24 hours before plating. Colony-forming unit (CFU) counts of BT ancestor and the V1 variant were determined by plating and counting on TSA Congo Red. Each condition included five biological replicates, with each replicate consisting of three technical replicates.

### Biofilm matrix staining with fluorescent lectins

Matrix glycoconjugate characterization in the three species community was performed using 78 different fluorescently labeled lectins (using FITC, AlexaFluor488, or Fluorescein) in combination with confocal laser scanning microscopy (TCS SP5X, Leica Germany, controlled by the software program LAS-AF vers. 2.41) as described previously by Neu and Kuhlicke [45, 85]. The lectins and fluorescent conjugates used are listed in Supplementary Table 5. Briefly, biofilms were grown on polycarbonate chips for 24 h, followed by washing three times with 1x PBS, as described above. Staining solutions containing fluorescent lectins were prepared at a concentration of 100 µg/mL (in filter-sterilized ultrapure water). 100 µL was then added to a biofilm on a polycarbonate chip, followed by incubation for 30 min in the dark. The excess staining solution was washed twice with dH_2_O before adding SYTO60 to the sample, prior to confocal imaging. Excitation of FITC, AlexaFluor488, and Fluorescein was performed at 488 nm, and the emission signals were recorded from 500–580 nm. Excitation of SYTO60 was at 652 nm, and emission was recorded from 665-750 nm. For lectins conjugated to Rhodamine, excitation was done at 525 nm, and emission was recorded from 570-650 nm. Image data sets were recorded using 63x NA 0.9 and 63x NA 1.2 objective lenses with a step size of 0.5 µm.

Based on the screening, three lectins, *Aleuria aurantia* Lectin (AAL), Wheat Germ Agglutinin (WGA), and *Ricinus communis* Agglutinin (RCA), were chosen for further testing. Here, monospecies and dual species biofilms of PB and BT ancestral or variant strains were used to better assess the effects of BT genotype and co-cultivation on AAL lectin binding. Images of AAL-lectin-stained biofilms of BT,variants V1, V2, V3 and V6 strains were analyzed using the biofilm image analysis software suite BiofilmQ [86]. Biofilm volume (biovolume) was calculated for each fluorescent channel: lectin (FITC channel) and biomass (SYTO60 channel) with a minimum of 6 images per sample (3 biological replicates and 2 images per replicate). Moreover, biovolume ratios (AAL-lectin-stained biovolume divided by the total biovolume) were calculated per image.

### Biofilm matrix proteome measurements

Mono-cultures and co-culture biofilms were grown for 24 h in TSB medium on PC chips as described above, followed by washing 3 times in 1x PBS to remove planktonic cells and biofilm pellicle. Five biological replicates per condition were processed, each composed of five PC chips that were combined to extract enough biofilm matrix material for proteomics analysis (technical replicates). The biofilm was then resuspended in 5 mL of 1x PBS by shaking at 1000 rpm for 10 min. The biofilm matrix was extracted from biofilms grown in either mono-culture or co-culture, resulting in a mixture of matrix components from both species, by the formaldehyde and NaOH extraction method [87, 88]. Briefly, 0.006 V formaldehyde 37 % (30 μl) was added to the biofilm samples and incubated for 1 h at 4 °C while mixing, followed by addition of 0.4 V NaOH 1 N (2 mL) and incubation for 3 h at 4 °C while mixing. Samples were then dialyzed using 3500 MWCO 22 mm SnakeSkin™ Dialysis Tubing (ThermoFisher Scientific) against 2.5 l milliQ water, overnight at 4 °C with stirring. Protein content in dialyzed samples was quantified using the Bio-Rad Protein assay, a modified Bradford assay [89], before freezing with liquid nitrogen. Finally, the extracted biofilm matrix was lyophilized using an ALPHA 1-4 LDplus freeze dryer (Christ, VWR) and kept at −80 °C until sample preparation for proteomics.

The proteomics workflow included sample preparation, trypsin digestion, mass spectrometry, and data analysis conducted as described in Herschend et al. [75] with some modifications.

### Matrix proteomics

Freeze-dried samples were resuspended in 100 µL lysis buffer (6M Guanidinium Hydrochloride, 10mM TCEP, 40mM CAA, 50mM HEPES pH8.5) and sonicated for 2 minutes in a sonication bath (Ultrasonic cleaner, VWR) at level 7-8. Samples were boiled for 5 min at 95 °C, spun down 10 min at 10000 rpm and 4 °C and supernatants transferred to Lo-bind microtiter plate. Protein content was quantified with Pierce™ Rapid Gold BCA protein assay kit (Thermo Scientific) and 10 μg protein material was used for digestion. The samples were three-fold diluted in digestion buffer (50 mM HEPES Ph 8.5 and 10% acetonitrile) containing LysC (1:50 enzyme:protein ratio), briefly vortexed and incubated for 1 hour at 37 °C. Then samples were diluted 10x in digestion with trypsin (1:100 trypsin-to-protein ratio), briefly vortexed and incubated at 37 °C temperature overnight. Inactivation of trypsin was achieved by adding trifluoroacetic acid (TFA) to 2% and debris was removed by centrifugation (x g, 1 min 4 °C). Acidified peptides were loaded onto a SOLAµ HRP 96-well plate for desalting, which was previously activated with 100% methanol, followed by 80% acetonitrile (AcN) complemented with 0.1% TFA and centrifugation at 1500 rpm for 1 min. SOLAµ HRP 96-well plate was then washed twice with 0.3% and 1% TFA and samples loaded and washed twice using 200 μL 0.1% TFA before being eluted with 2 x 30 µl 40% AcN complemented with 0.1% TFA. AcN was removed from eluted samples in a SpeedVac for at least 1 hour and 60 °C. Dry peptide samples were resuspended in 12 µL A*iRT solution, sonicated for 60 seconds and quantified using a DS-11 drop spectrophotometer. Desalted and purified peptide samples were kept at −80 °C until subjected to analysis by mass spectrometry.

### Mass Spectrometry analysis

For each sample, peptides were loaded onto a 2cm C18 trap column (ThermoFisher 164705), connected in-line to a 15cm C18 reverse-phase analytical column (Thermo EasySpray ES803) using 100% Buffer A (0.1% Formic acid in water) at 750bar, using the Thermo EasyLC 1000 HPLC system, and the column oven operating at 45 °C. Peptides were eluted over a 190 minute gradient ranging from 10 to 60% of 80% acetonitrile, 0.1% formic acid at 250 nl/min, and the Q-Exactive instrument (Thermo Fisher Scientific) was run in a DD-MS2 top10 method. Full MS spectra were collected at a resolution of 70,000 with an AGC target of 3×10^6^ or maximum injection time of 20 ms and a scan range of 300–1750 m/z. The MS2 spectra were obtained at a resolution of 17,500, with an AGC target value of 1×10^6^ or maximum injection time of 60 ms, a normalised collision energy of 25 and an intensity threshold of 1.7e^4^. Dynamic exclusion was set to 60 s, and ions with a charge state <2 or unknown were excluded. MS performance was verified for consistency by running complex cell lysate quality control standards, and chromatography was monitored to check for reproducibility.

### Proteomics data analysis

Peptides identified by mass spectrometry, representing a mixture from both species, were mapped to the reference proteomes of *Bacillus thuringiensis* and *Pseudomonas brenneri* using MaxQuant v1.6.10.43 [90] with the built-in Andromeda search engine [91]. A target decoy search that allowed for a maximum of 1% FDR on both peptide and protein level and a minimum length of 7 amino acids per peptide was performed. Quantification was performed using the label-free quantification (LFQ) algorithm [92] in MaxQuant with a minimum ratio count of 2 and applying the “match between runs” function. The Andromeda search engine was supplemented with the reference proteome of *Bacillus thuringiensis,* and/or *Pseudomonas brenneri*. The genomes had been prepared during a previous study [29] and are available online (NCBI) with the following IDs: IDs JARTVI000000000 (*B. thuringiensis*) and CP122540.1 (*P. brenneri*).

Statistical analysis of proteomics data was performed with the Perseus software v1.6.15.0 [93] and differentially expressed proteins were identified using a modified version of Welch’s t-test with an S0 parameter of 0.5 [94] and a permutation-based FDR cut-off of 0.05 and valid values in at least 60% of the samples in both sample groups, depending on the comparison made. In this software, pairwise comparisons were performed for each species using the software Perseus: for *Bacillus thuringiensis*, monoculture versus co-culture (for both ancestral and variant genotypes); and for *Pseudomonas brenneri*, monoculture versus co-culture (including ancestral, V1, and V6 genotypes).

Prediction of subcellular localization of detected proteins was done using subcellular localization prediction software PSORTb version 3.0.3 (https://www.psort.org/psortb/) [95]. Similarly, prediction signal peptides and the location of their cleavage sites in the detected proteins was done using SignalP 6.0 (https://services.healthtech.dtu.dk/service.php?SignalP-6.0) [96].

### Statistical analysis and visualization

All analyses and plotting were done in RStudio v4.1.1 [97] and package “ggplot2” v3.4.2. All datasets were tested for normality and equal variance using the Shapiro-Wilk (base R) and Levene’s (package “car” v3.1.2) tests, respectively. Most datasets were not normally distributed and thereby a Kruskal-Wallis test (base R) was used when comparing more than two groups, unless otherwise specified. A general expression was kruskal.test(log10_CFUmL∼Combination), however the quantitative variable and categorical variables changed depending on the dataset. A Dunńs post-hoc test was performed using package “FSA” v0.9.4 to consider multiple comparisons when the Kruskal-Wallis test was significant, with a Bonferroni correction. When comparing only two groups, a Wilcoxońs rank test t-test was applied, usually comparing ancestor and variant phenotype/genotype (i.e. wilcox.test (CFU∼Phenotype)). A *p*-value of 0.05 was used as a threshold for statistical significance (*p* < 0.05). Compact letter display for multiple comparisons was calculated using the R package “rcompanion” v2.4.35.

## Author contributions

CIA is first author. Conceptualization and planning: HLR. Execution of experiments: CIA, HLR. SZM, ISK. Data analysis: CIA, HLR and LM. Manuscript preparation: CIA, HLR and MB, in consultation with JH, HJ, VSC, KD, TRN.

## Acknowledgments

The authors thank Sara Olsen and Anette Løth for technical assistance, the DTU Proteomics core facility for technical assistance in proteomics sample preparation. Support by Ute Kuhlicke with lectin analysis and confocal laser scanning microscopy is highly appreciated. We thank the Villum Foundation for grants to MB (no. 10098) and HR (no. 34434) and the US Office of Naval Research, for grant N629091812174 to MB, as well as the Swiss National Science Foundation Consolidator Grant TMCG-3_213801 to KD, and the German Bundesministerium für Bildung und Forschung grant TARGET-Biofilms 16GW0245 to KD, funding this work.

## Data availability statement

*B. thuringiensis* and *P. brenneri* genomes are available online at NCBI (https://www.ncbi.nlm.nih.gov/genome/) with the following IDs: JARTVI000000000 and CP122540.1, respectively. The mass spectrometry proteomics raw data and results from analysis by MaxQuant have been deposited to the ProteomeXchange Consortium via the PRIDE [98] partner repository with the dataset identifier PXD046156 (https://www.ebi.ac.uk/pride/archive). The data is currently private and can only be accessed with a single reviewer account that has been created. The data will be publicly available when accepted for publication. For reviewer access, use the username and password kkAMO14X.

---

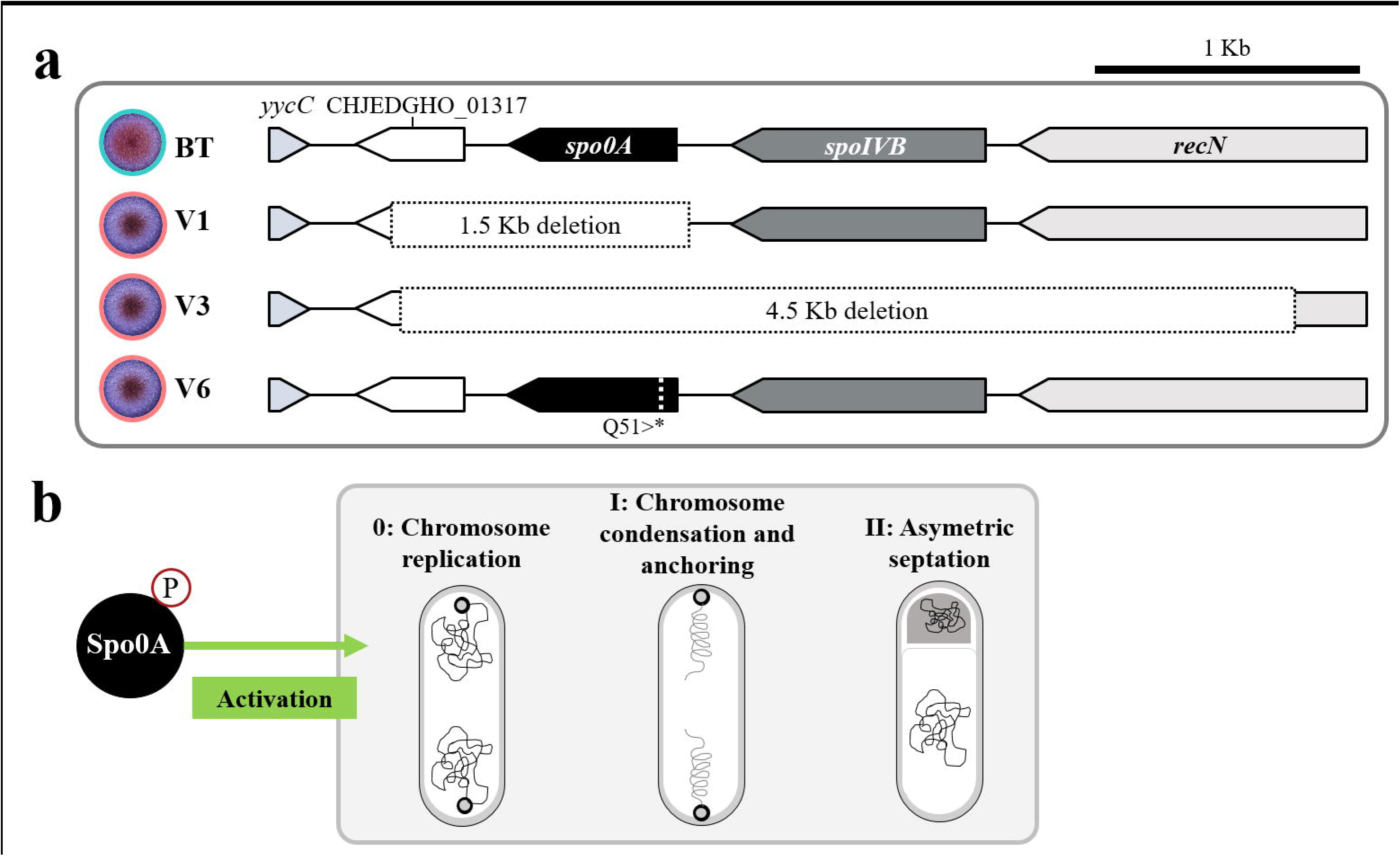

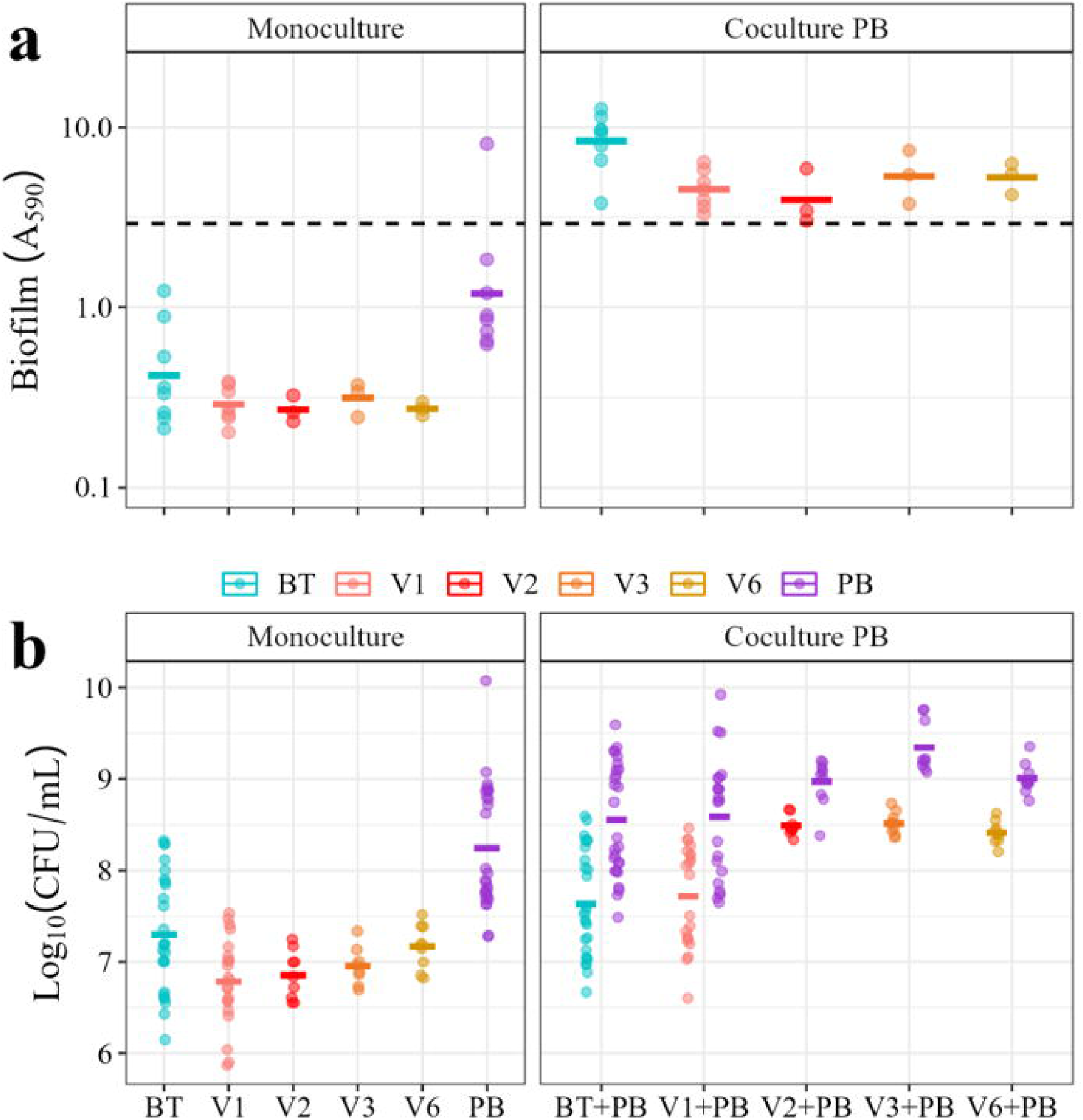

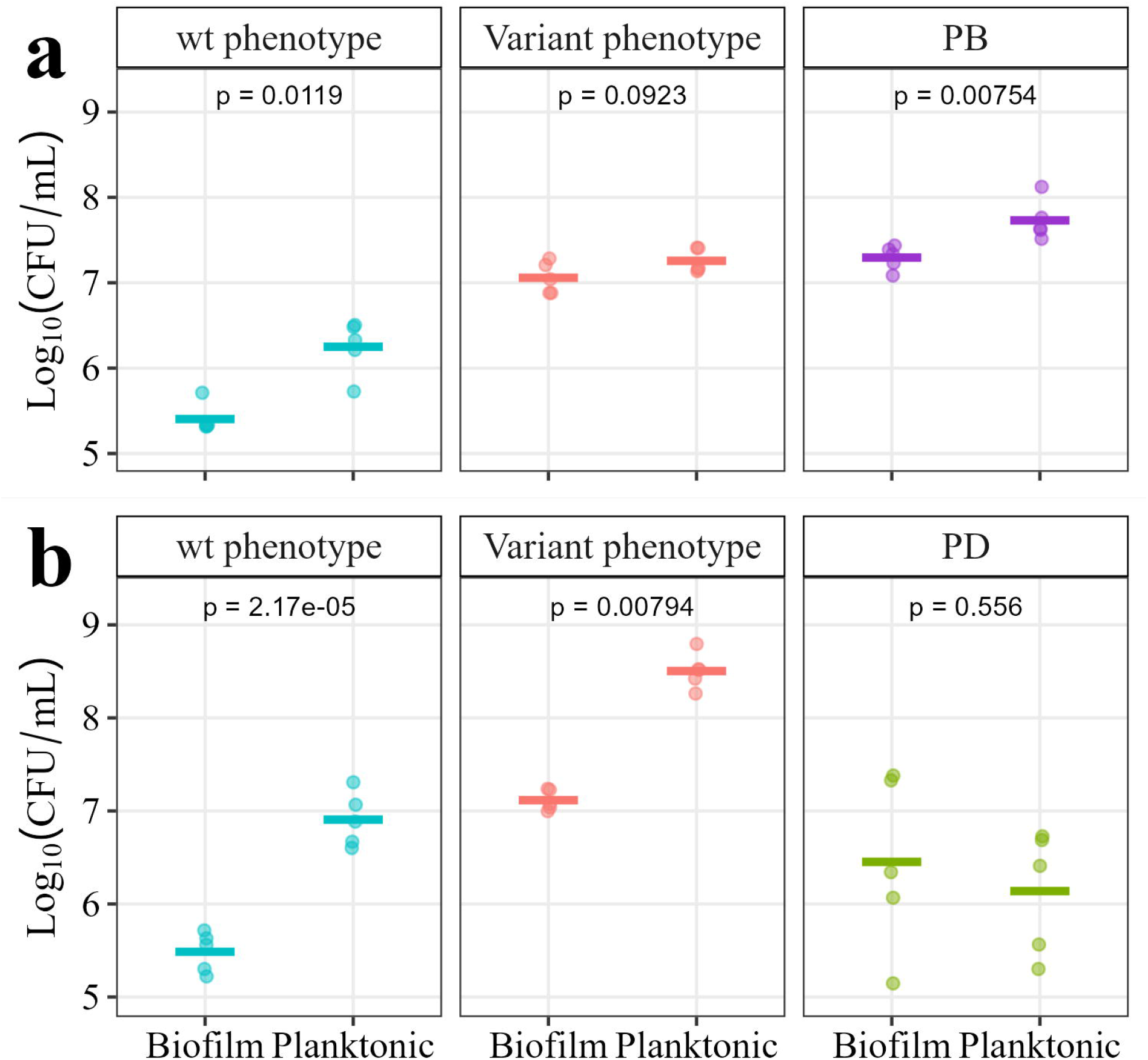

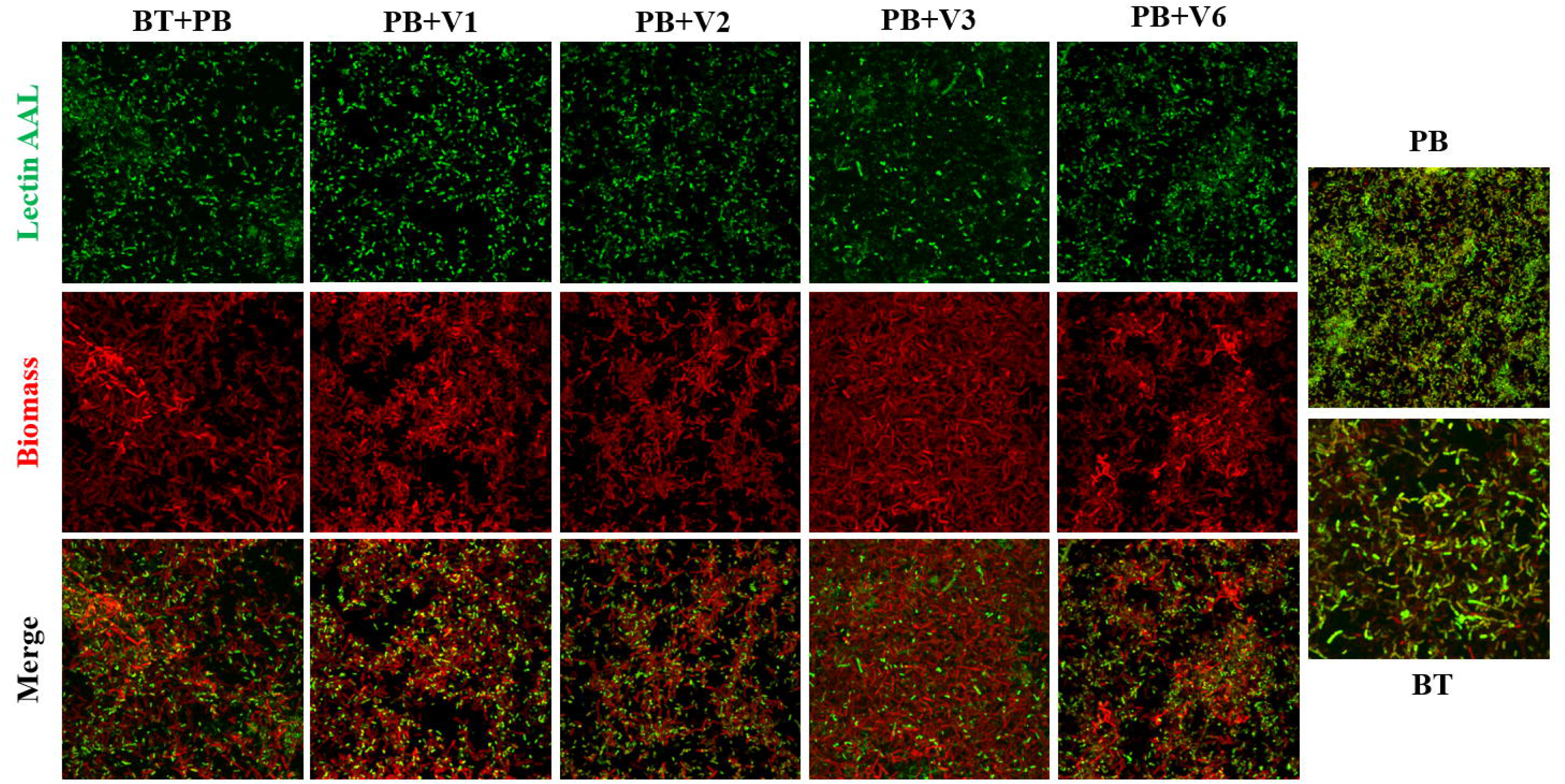

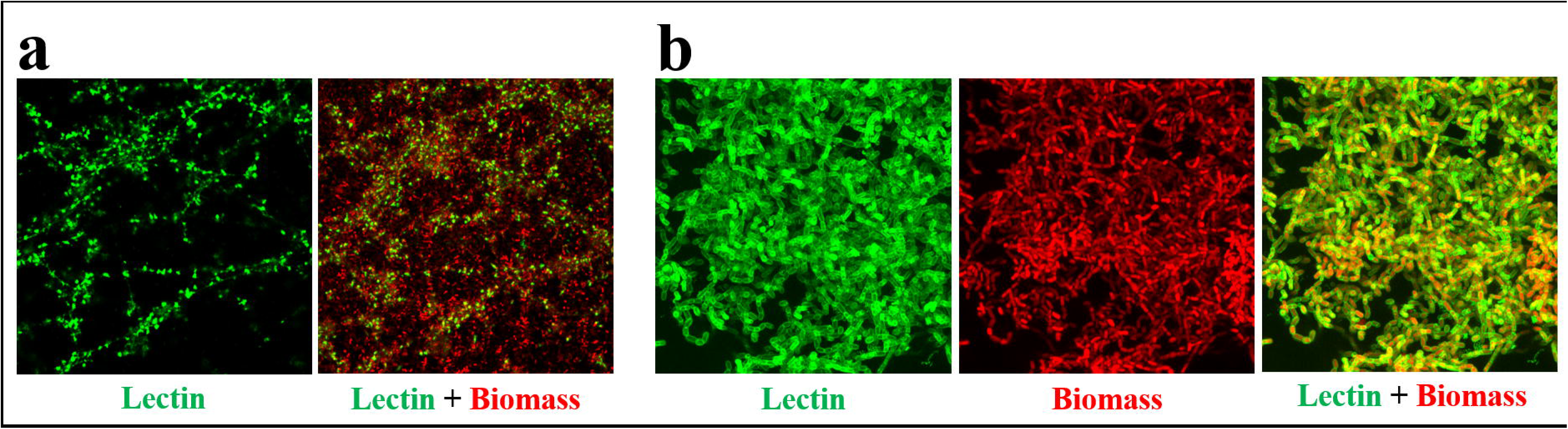

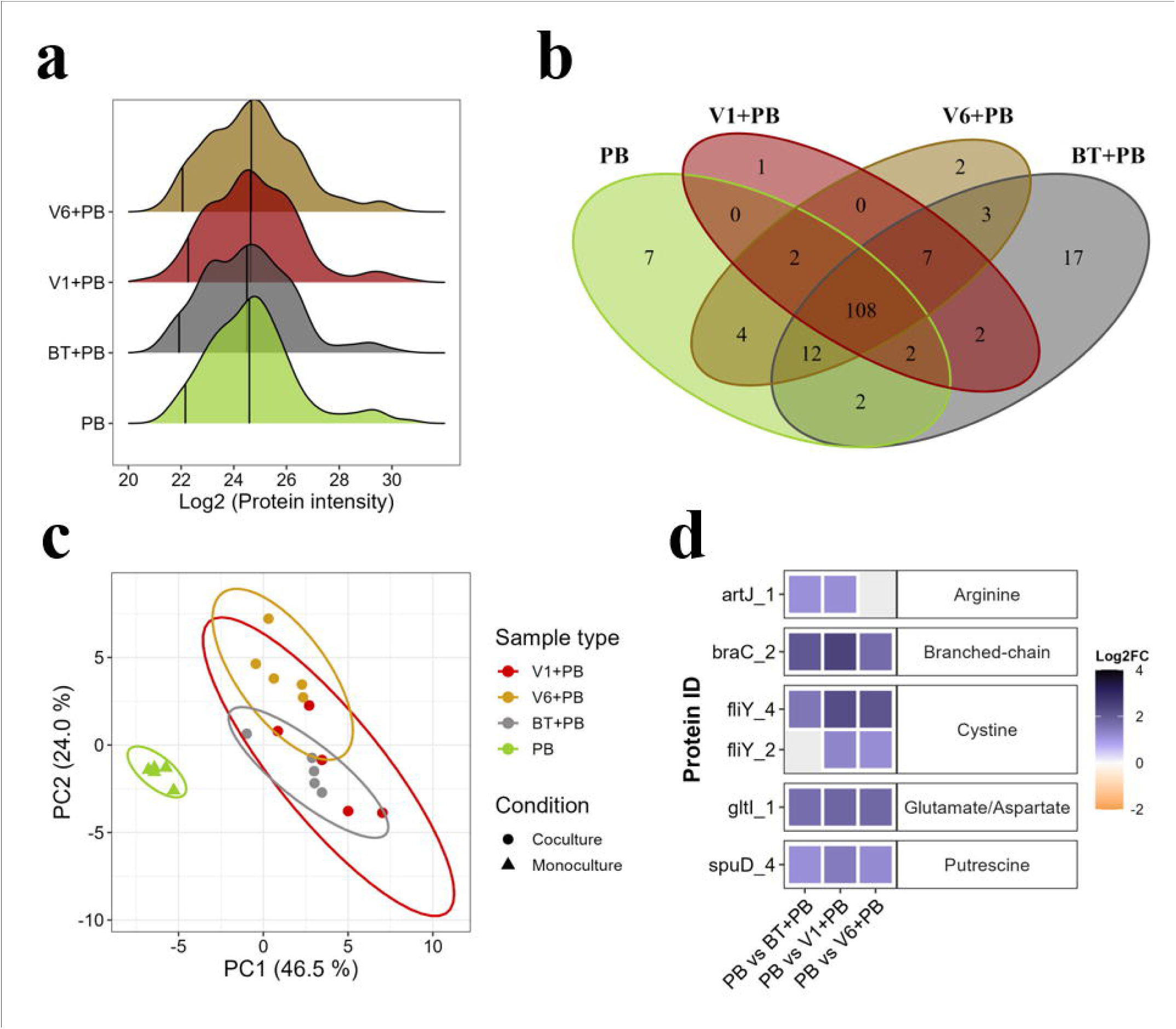

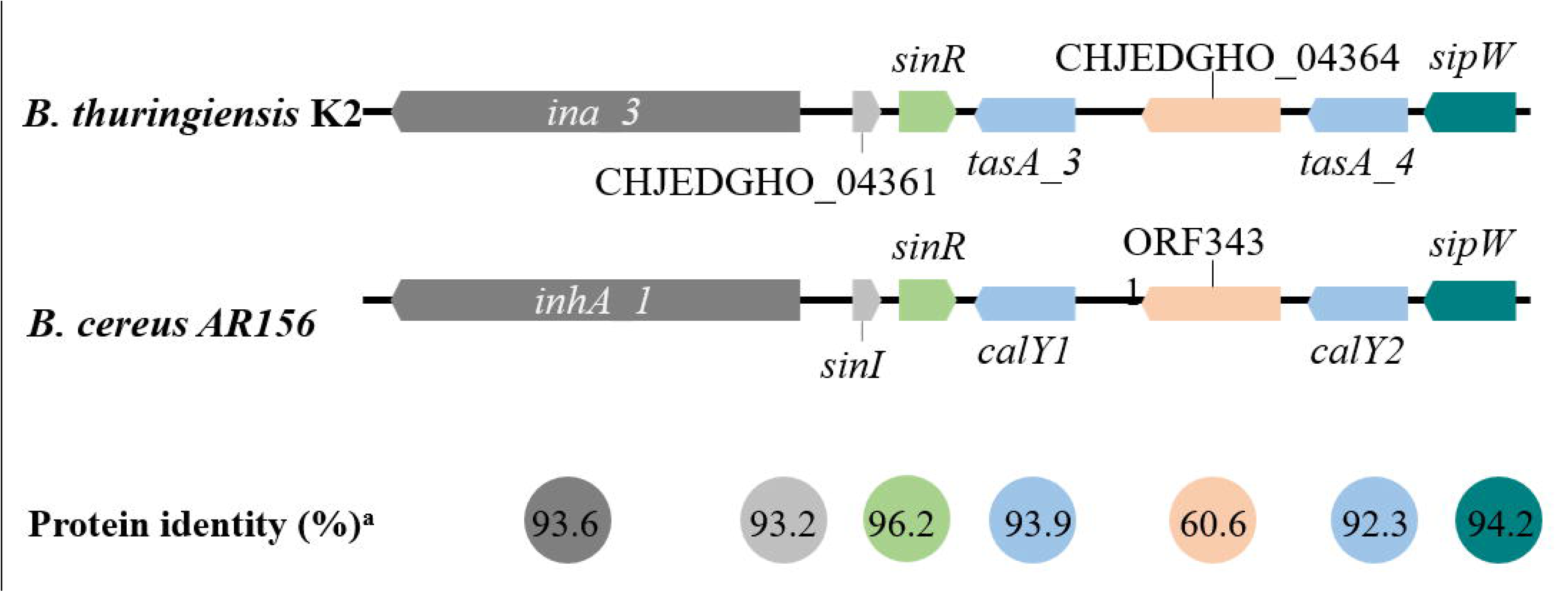

