## Supplementary material for "Evolution of genotypic and phenotypic diversity in multispecies biofilms"

[2]Present address: Functional Foods, Novozymes A/S, 2800, Lyngby, Denmark

[6]Pressent address: Section of Microbiology and Fermentation. Department of Food Science, University of Copenhagen. Rolighedsvej 26, 1958 Frederiksberg C, Denmark.

**Keywords**: multispecies, bacterial interactions, biofilm, matrix, adaptive evolution, sporulation

**The supplementary file includes:**

Supplementary figures 1-7

Supplementary tables 1-13 are available in supplementary file “Amador et al. 2024 Supplementary tables.xlsx”.

#### Supplementary figures

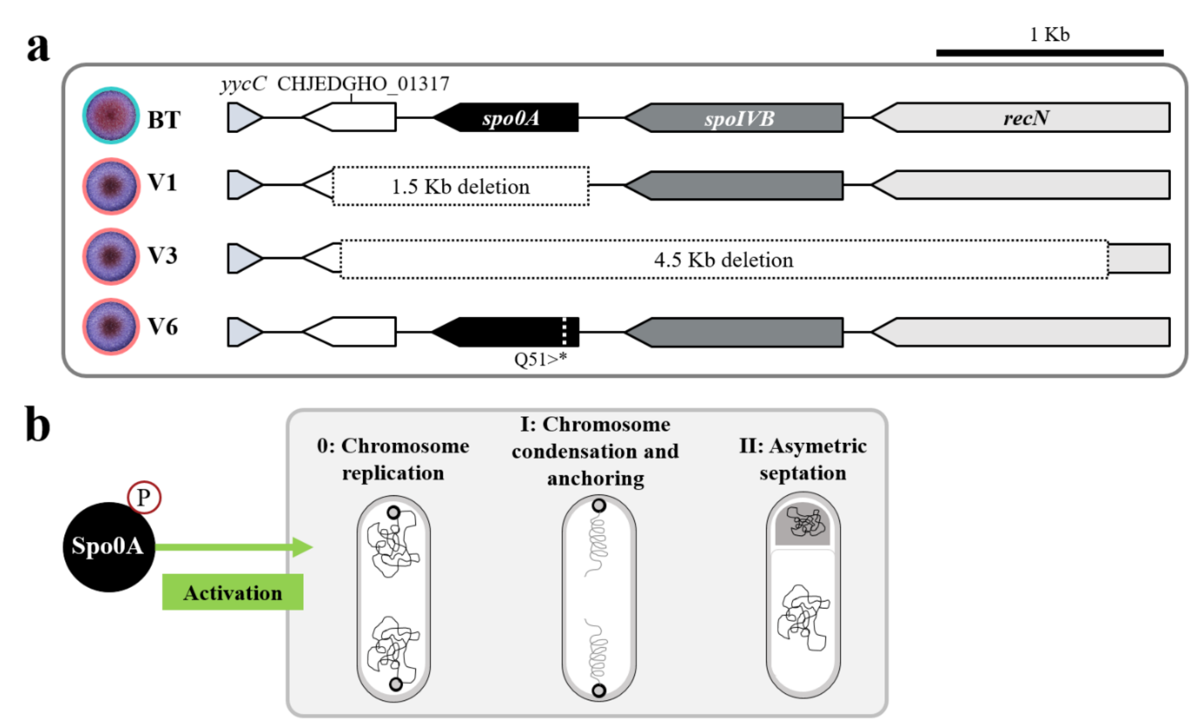


**Supplementary Figure 1. Mutations in *B. thuringiensis* variants compared to the ancestral strain and its effect on sporulation pathway. (a)** Mutations found in the selected variants (BT Vx, circled pink) compared to their ancestor (BT, circled blue). Dashed boxes indicate large deletions in variants V1 and V3. Q51>* in variant V6 indicates a change in amino acid 51 of the Spo0A amino acid sequence from glutamine (Q) to a stop codon (*). **(b)** Role of SpoA in activation of the sporulation pathway in *Bacillus* sp. Phosphorylated Spo0A activates transcription of genes involved in sporulation stages 0, I, and II [1, 2].

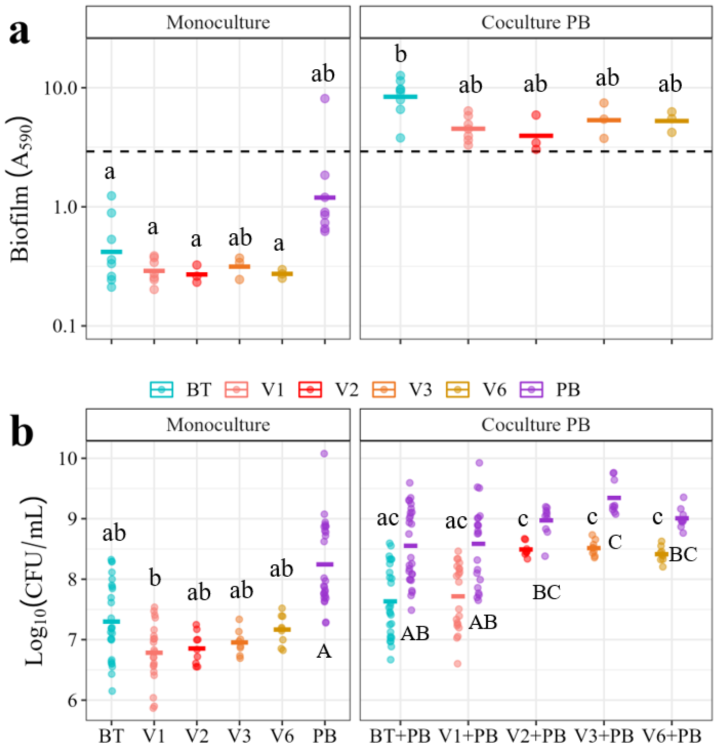


**Supplementary Figure 2. Biofilm formation of *B. thuringiensis* ancestral and variant strains in mono-culture or co-culture with *P. brenneri* after 24 hours.** Each color indicates a different strain. BT: *B. thuringiensis* ancestor; V1: V1 variant; V2: V2 variant, V3: V3 variant; V6: V6 variant; PB: *P. brenneri*. Each panel represents biofilm after 24 hours measured in either mono-culture or co-culture with *P. brenneri* (Coculture PB). **(a)** Total biofilm in TSB measured as crystal violet stained biomass ($A_{590}$). Data is based on ≥ 9 biological replicates, represented as individual points, and 3 technical replicates. Mean values are represented by horizontal crossbars. Dissimilar letters denote significant difference in multiple comparison per panel ($p < 0.05$, Dunn´s post-hoc test). **(b)** Biofilm measured as CFU adhered per mL in TSB after 24-hour incubation. Data based on ≥ 9 biological replicates, indicated by individual points, and ≥ 3 technical replicates. Dissimilar letters denote significant difference in multiple comparisons per strain, *P. brenneri* in capital letters. ($p < 0.05$, Dunn´s post-hoc test).

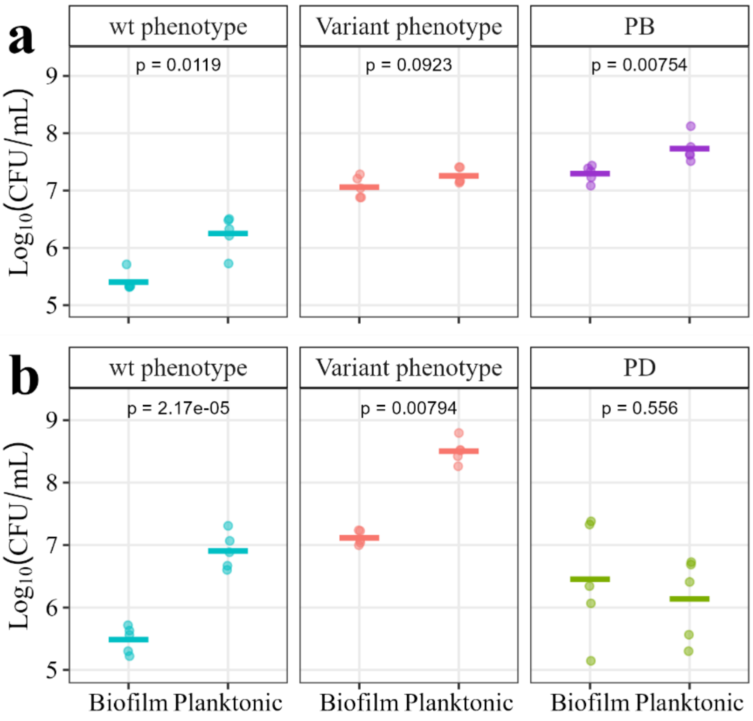


**Supplementary Figure 3**. **Competition of *B. thuringiensis* ancestral and V1 variant strains under biofilm or planktonic settings in the presence of only one *Pseudomonas* spp.** Planktonic samples were cultivated for 24 h in TSB, diluted 1:1000 into fresh medium, and incubated for another 24 h before plating. Biofilm samples were cultured for 24 h, washed thrice, transferred to fresh media, and regrown for 24 h before plating. Plots show mean CFU counts (horizontal crossbars) of wild-type and variant colony phenotypes ($Log_{10}$(CFU/mL)) per setting and combination. Data represent five biological replicates (points), each with three technical replicates. P-values ($p$) indicate significance of biofilm vs. planktonic CFUs per strain ($p < 0.05$, Two-sample t-test). (**a**) Competition of BT ancestor and V1 variant in the presence of PB (BT+V1+PB). PB = *P. brenneri.* (**b**) Competition of BT ancestor and V1 variant in the presence of PD (BT+V1+PD). PD = *P. defluvii.*

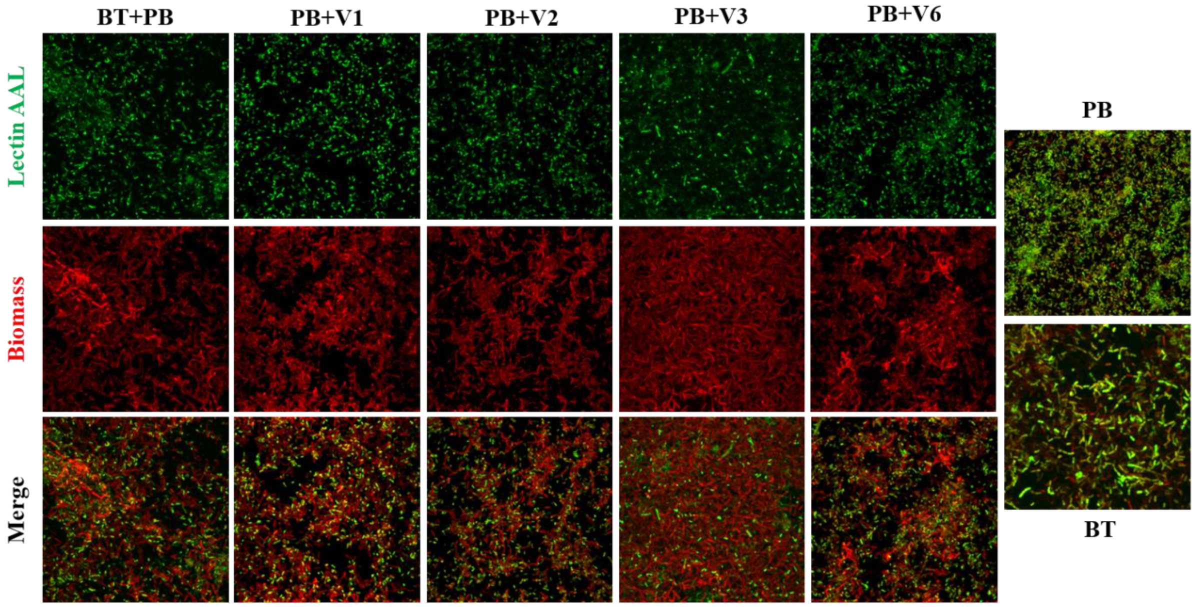


**Supplementary Figure 4. Co-culture biofilms of *B. thuringiensis* ancestral and variant strains with *P. brenneri*, stained with AAL lectin and SYTO60.** BT = *B. thuringiensis* ancestral strain, V1 = BT variant 1, V2, BT variant 2, V3 = BT variant 3, V6 = BT variant 6, PB = *P. brenneri*. Maximum intensity projection (MIP) images of Z-stacks recorded by confocal laser scanning microscopy for 24-hour biofilms. Lectin AAL: fluorescent images of biofilms stained with the fluorescent lectin AAL-FITC. Biomass: fluorescent images of biofilms stained with the cell biomass stain SYTO60. Merge: combined images of biomass and lectin stains. BT: Images represent 124 μm x 124 μm.

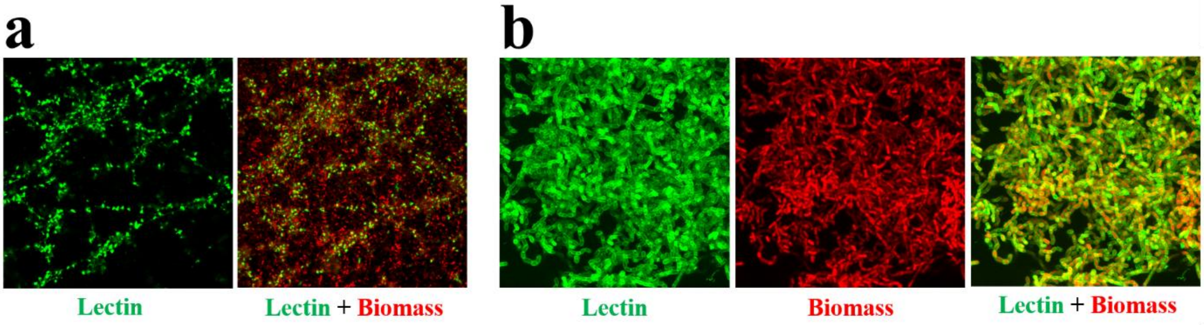


**Supplementary Figure 5. RCA and WGA lectin binding to *P. brenneri* and *B. thuringiensis* biofilms.** Maximum intensity projection (MIP) images of Z-stacks recorded by confocal laser scanning microscopy of 24-hour biofilms. Lectin binding is shown in green and SYTO60 cell biomass stain in red. Images represent 124 μm x 124 μm. **(a)** RCA-FITC lectin binding to *P. brenneri* biofilms. **(b)** WGA-FITC lectin binding to *B. thuringiensis* ancestor biofilm.

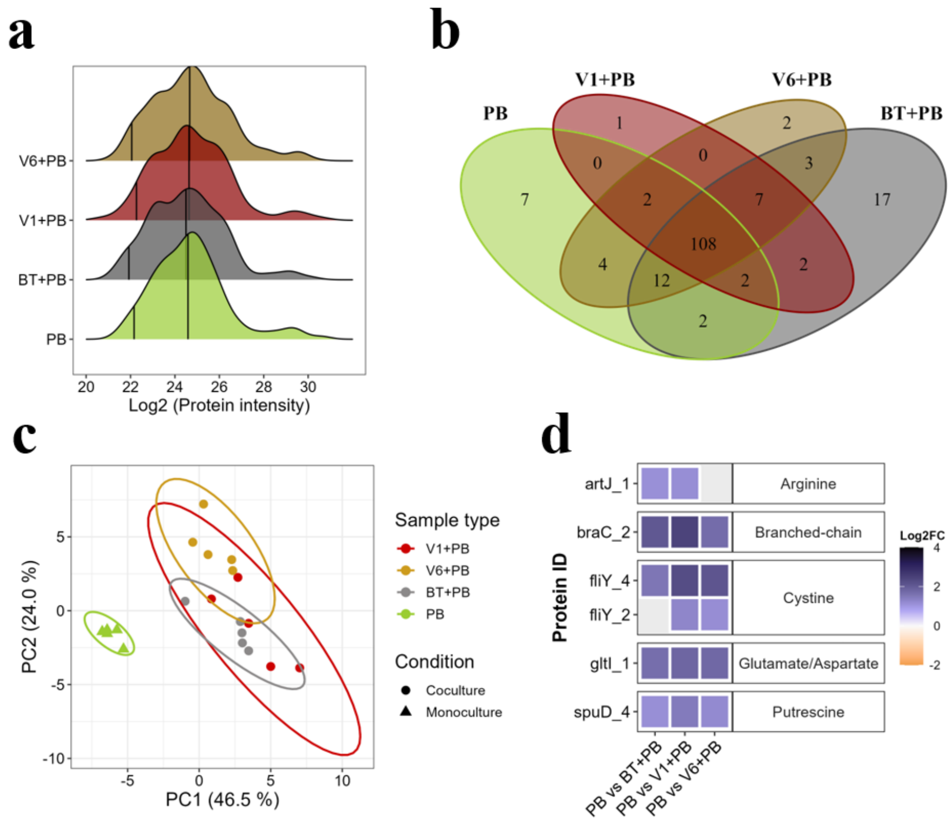


**Supplementary Figure 6. Biofilm proteomes of *P. brenneri* in mono-culture or co-culture with *B. thuringiensis* ancestral and variant strains.** PB (green) = *P. brenneri* mono-culture; BT+PB (grey): co-culture with *B. thuringiensis* ancestral strain; V1+PB (dark red): co-culture with V1 variant; V6+PB (dark yellow): co-culture with V6 variant. **(a)** Density plots of Log2 transformed protein intensity for mono-culture vs. co-culture samples representing five biological replicates. Percentile 0.05 is represented as the leftmost vertical line, while middle lines denote protein intensity means for each sample excluding NA values (no protein detected). **(b)** Venn diagram of *P. brenneri* mono- and co-culture proteomes, total proteins detected = 169, considering proteins detected in ≥ 60 % of each sample (3/5 replicates). PB mono-culture, n = 137 proteins; V1+PB, n = 122 proteins; V6+PB, n = 138 proteins; BT+PB, n = 153 proteins. **(c)** Probabilistic PCA plot of mono-culture and co-culture proteomes based on 5 biological replicates per sample. Color indicates the sample group while shape distinguishes culture conditions: co-culture (circles) or mono-culture (triangles). Ellipses indicate sample group clustering based on the principal components. **(d)** Differential abundance of proteins within "Amino acid transport and metabolism" category (COG classification [3]) in mono-culture vs. co-culture with *B. thuringiensis* ancestor or variants, expressed as $Log_2$ fold change ($Log_2FC$). Negative values indicate higher protein abundance in co-culture while positive values higher abundance in mono-culture.

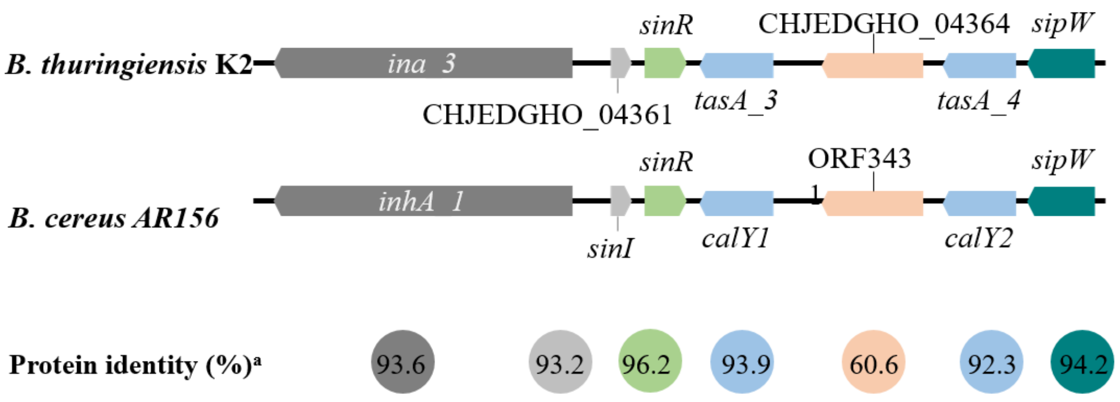


**Supplementary Figure 7. Genomic arrangement of the *sinI–sinR* region in *Bacillus thuringiensis* K2 and *Bacillus cereus* AR156.** The SinI-SinR regulatory circuit in *B. cereus* AR156 was previously described by Xu et al. [5]. The corresponding genomic region from *B. cereus* AR156 (GenBank accession number CP015589.1) was retrieved and aligned to the *Bacillus thuringiensis* K2 genome [4] using BLASTp (https://blast.ncbi.nlm.nih.gov/Blast.cgi).[a] The percentage identity between proteins in the sinI–sinR region of *B. cereus* AR156 and *B. thuringiensis* K2 was calculated using Clustal Omega (https://www.ebi.ac.uk/jdispatcher/msa/clustalo).
